# Selective LXR agonist, DMHCA, corrects the retina-bone marrow axis in type 2 diabetes

**DOI:** 10.1101/2020.02.11.942292

**Authors:** Cristiano P. Vieira, Seth D. Fortmann, Masroor Hossain, Ana Leda Longhini, Sandra S. Hammer, Bright Asare-Bediako, David K. Crossman, Micheli S. Sielski, Yvonne Adu-Agyeiwaah, Mariana Dupont, Jason L. Floyd, Sergio Li Calzi, Todd Lydic, Robert S Welner, Gary J. Blanchard, Julia V. Busik, Maria B. Grant

## Abstract

In diabetic dyslipidemia, cholesterol accumulates in the plasma membrane, decreasing fluidity and thereby suppressing the ability of cells to transduce ligand-activated signaling pathways. Liver X receptors (LXRs) are the main cellular mechanism by which intracellular cholesterol is regulated and play important roles in inflammation and disease pathogenesis. N,N-dimethyl-3β-hydroxy-cholenamide (DMHCA), a selective LXR agonist, specifically activates the cholesterol efflux arm of the LXR pathway without stimulating triglyceride synthesis. Thus, DMHCA possesses superior clinical potential as a cholesterol lowering agent than current LXR pan-agonist. In this study, we use a multi-systems approach to understand the effects and molecular mechanisms of DMHCA treatment in type 2 diabetic db/db mice and human -derived circulating angiogenic cells (CACs), which are vascular reparative cells. We find that DMHCA is sufficient to correct the retina-bone marrow (BM) axis in diabetes, thereby restoring retinal structure, function, and cholesterol homeostasis, rejuvenating membrane fluidity in circulating vascular reparative cells, hampering systemic inflammation, and correcting BM dysfunction. Using single-cell RNA-seq on lineage^-^sca1^+^cKit^+^ (LSK) hematopoietic stem cells (HSCs) from untreated and DMHCA-treated diabetic mice, we provide novel insights into hematopoiesis and reveal DMHCA’s mechanism of action in correcting diabetic HSCs by reducing myeloidosis and increasing CACs and erythrocyte progenitors. Taken together, these findings demonstrate the broad and pleiotropic effects of DMHCA treatment, which has exciting potential to correct the retina-BM axis in diabetic subjects.

## Introduction

The landmark ACCORD Eye study demonstrated that, in type 2 diabetic (T2D) individuals with dyslipidemia, tight glycemic control supplemented with fenofibrate/statin combination significantly reduced the progression of diabetic retinopathy (DR) compared to statin supplement alone (1). Subgroup analysis of the dyslipidemia cohort revealed that elevated LDL cholesterol was the only individual lipid measurement that was significantly associated with worse DR progression. These data establish diabetic dyslipidemia, and hypercholesterolemia specifically, as risk factors for DR and support the notion that therapies targeting lipid metabolism are clinically efficacious in T2D.

Serum cholesterol, which reflects the cholesterol exchange between tissues, is the clinical measurement used to estimate an individual’s total cholesterol level. However, the overwhelming majority of cholesterol is distributed in the cell membranes of peripheral tissues where it accounts for 30-50% of the plasma membrane molar ratio (2). Statins, the first line treatment for hypercholesterolemia, predominantly target cholesterol biosynthesis, and thereby decrease circulating LDL (3). However, statins have a lesser effect on the efflux of intracellular cholesterol in peripheral tissues such as the retina (4, 5). In diabetic dyslipidemia, cholesterol accumulation leads to changes in membrane fluidity, inflammation and disease pathogenesis. (6, 7). Increase in cholesterol could have pleotropic and cell-specific effects. We have previously demonstrated that in human retinal endothelial cells (HREC), increase in membrane cholesterol leads to stabilizations of membrane microdomains, such as lipid rafts, which promote cytokine receptor clustering leading to increased intracellular second messenger signaling and amplification of inflammatory cytokine signaling (8, 9). In addition, cholesterol can affect membrane fluidity that is of particular importance for bone marrow-derived cell trafficking and mobility.

Liver X receptors (LXR) are the main cellular mechanism by which intracellular cholesterol is regulated. These nuclear receptors transcriptionally regulate genes involved in lipid metabolism to homeostatically balance the endogenous biosynthesis, dietary uptake, metabolism, and elimination of lipids (10). LXR activation is induced by elevated intracellular cholesterol and stimulates cholesterol removal through reverse cholesterol transport (11, 12). In addition, LXR activation maintains the composition and physical properties of the cell membrane through the coupled regulation of phospholipid and cholesterol metabolism (13). Activation of LXR is signaled through direct binding of endogenous lipid ligands, such as oxysterols and other cholesterol derivatives, as well as intermediate precursors in the cholesterol biosynthesis pathway (10, 11). Synthetic chemical agonists of LXR have been developed for therapeutic intervention, and while they have proven efficacious in diabetic animal models (14–18), their undesirable adverse effect profile, including hypertriglyceridemia and hepatic steatosis, has hampered clinical development (19, 20).

N,N-dimethyl-3β-hydroxy-cholenamide (DMHCA) is a synthetic oxysterol that induces gene-specific modulation of LXR (21). Mechanistically, DMHCA indirectly activates LXR through the inhibition of desmosterol reduction, the final step in the predominant cholesterol biosynthesis pathway, leading to the accumulation of the potent LXR agonist desmosterol (22). During endogenous LXR activation, fatty acid biosynthesis is stimulated through the transcriptional induction of SREBP1c, leading to elevated triglyceride levels (23). Intriguingly, DMHCA selectively activates the cholesterol efflux arm of the LXR pathway, through the induction of ATP-binding cassette transporter (ABCA1), with minimal effect on SREBP1c compared to other LXR agonists, T0901317 and GW3965 (21–25). Thus, DMHCA has superior clinical potential as a cholesterol lowering agent because it lacks the undesirable adverse effect profile that plagued the first generation of LXR modulators, while retaining the ability to lower circulating LDL and restore peripheral cholesterol homeostasis (21, 23).

LXR signaling plays an important role in inflammation and disease (26). In the retina, LXR depletion causes retinal/optic nerve degeneration (27) and the formation of acellular capillaries (17) and retinal pigment epithelial changes (28), suggesting that LXR is required for normal retinal maintenance and its absence results in pathologies spanning the entire retina. Interestingly, in the diabetic retina, LXR expression is downregulated, and activation of LXR using a chemical agonist is sufficient to reduce gliosis and the formation of acellular capillaries (17, 18). LXR activation also displays potent anti-inflammatory effects, which are mediated, in part, by altering the composition of the plasma membrane (29). By selectively regulating the cholesterol content of specific membrane microdomains, LXR inhibits signaling through toll-like receptors (TLRs) 2, 4, and 9 (29). In diabetes, gut barrier dysfunction is an early event which increases circulating bacterial antigens, leading to enhanced activation of TLRs on endothelium and promoting chronic systemic inflammation (30). Thus, LXR agonists, such as DMHCA, have the added potential benefit of hampering widespread inflammation in diabetes.

Additional features of diabetes are vascular insufficiency and deficient wound healing. Circulating CD34^+^ vascular-reparative cells, also known as circulating angiogenic cells (CACs), play an important role in promoting vascular integrity and maintenance (31). These cells require a complex network of intercellular signaling to home to areas of injury and provide trophic support that promotes vascular repair. In diabetes, these cells are defective, low in number, and their levels correlate strongly with presence of microvascular complications, such as diabetic retinopathy (32, 33). Cell replacement treatments using nondiabetic-derived vascular-reparative cells have proven efficacious in DR mouse models (34), but drug treatments are needed that target and rejuvenate this population of circulating cells. In diabetic mice, LXR activation has been shown to restore the equivalent population of vascular reparative cells by enhancing their migration and suppressing oxidative stress and inflammatory gene expression (17). Moreover, in LXR double-knockout mice fed a high fat diet, circulating vascular reparative cells are dysfunctional, decreased in number, and show an increased cellular cholesterol content (35). Interestingly, these mice also demonstrated alterations in hematopoietic stem and progenitor cells (HS/PS), suggesting that LXR’s beneficial effects may extend to the hematopoietic stem cell (HSC) compartment (36). Together, these studies suggest that LXR modulators like DMHCA have the potential to prevent and treat diabetic complications spanning multiple tissues and cell types.

In this study, we use a multi-systems approach to understand the effects and molecular mechanisms of DMHCA treatment in T2D db/db mice and human CD34^+^ vascular reparative cells. Using lipidomics, single-cell membrane fluidity assays, flow cytometry, and single-cell RNA sequencing, we characterize the effects of diabetes and DMHCA treatment on the retina, circulating immune cells, and the BM.

## Results

### Systemic DMHCA treatment restores cholesterol homeostasis in the diabetic retina

In diabetes, dyslipidemia promotes the accumulation of intracellular sterols. To test whether selective LXR agonism, using systemic DMHCA treatment, is sufficient to reduce the levels of intracellular sterols in the retina, T2D db/db mice were treated with oral DMHCA for 6 months after the onset of diabetes. Liquid chromatography–mass spectrometry (LC-MS) was performed on lipid extracts from whole retina to quantify free (Figure 1A) and total sterols (Figure 1B; total sterols = free sterols + esterified sterols). In db/db diabetic mice compared to db/m heterozygous controls, the cholesterol content of the diabetic retina was roughly 1.5 magnitudes higher, consistent with diabetic dyslipidemia (Figure 1F). Since cholesterol accounts for the overwhelming majority of cellular sterols, the total sterol content was also increased by a similar degree.

**Figure 1.**
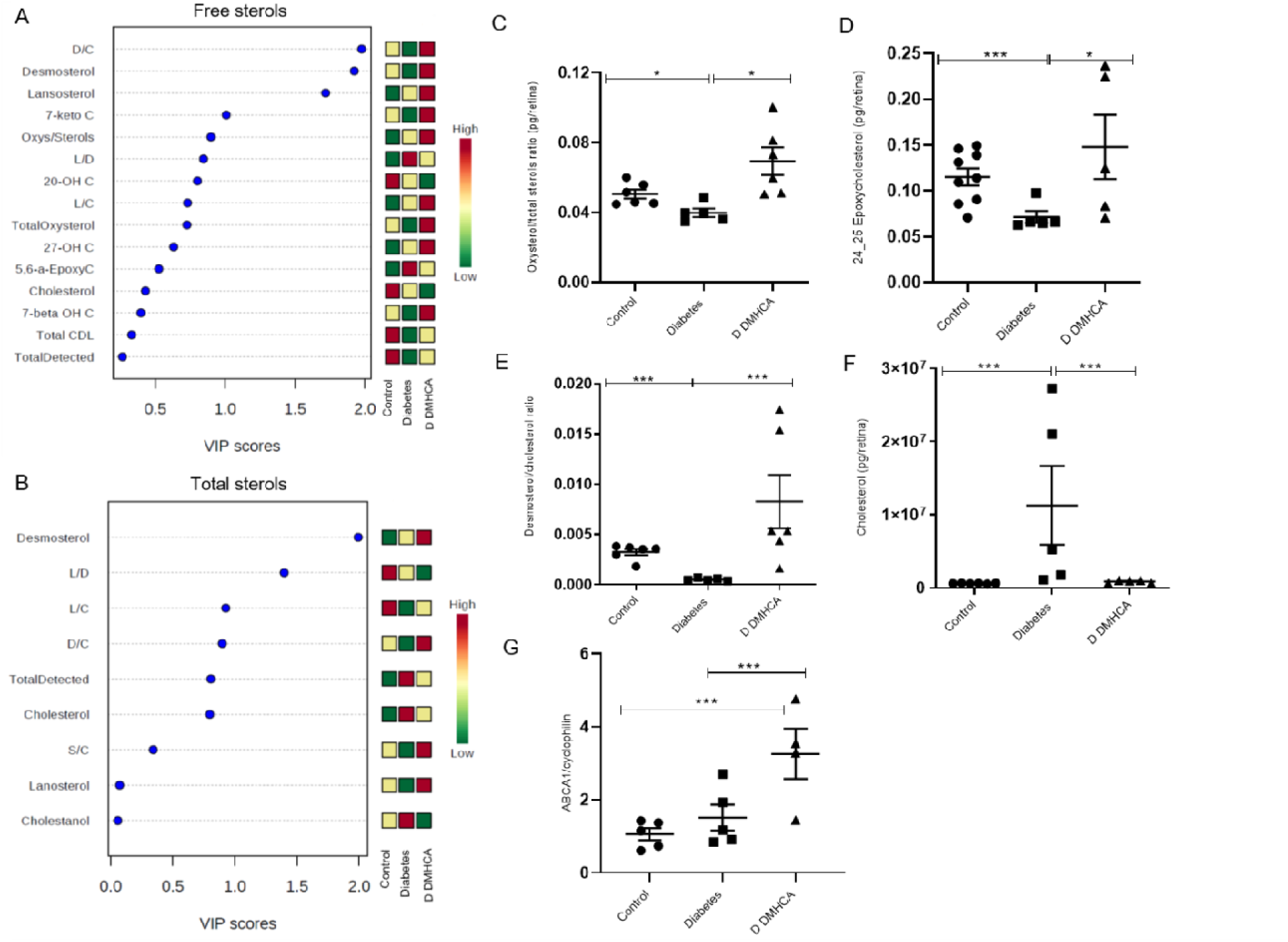
DMHCA restores cholesterol homeostasis in diabetic retina. Partial least squares discriminant analysis (PLS-DA) on LC-MS quantified retinal sterols from 9 month old mice. Total sterols = free sterols + esterified sterols. Colored boxes on the right indicate the relative concentrations of the corresponding metabolite for each group. CT = control (db/m); D = diabetes (untreated db/db); DD = DMHCA (treated db/db) (**A,B**). Quantification of total oxysterols/total sterols ratio(**C**), total 24_25 epoxycholesterol(**D**), total desmosterol/total cholesterol ratio(**E**), and total cholesterol(**F**). RT-qPCR on retinal ABCA1 mRNA expression(**G**). Data are the mean ± SEM. \**P* < 0.05; \*\**P* < 0.03 \*\*\**P* < 0.01 analyzed using unpaired 2-tailed Mann-Whitney test.

Interestingly, the correlation between free and total sterols for certain lipid species in the diabetic retina was inversely related. For example, free cholesterol in the diabetic retina was surprisingly decreased (Figure 1A), while esterified cholesterol was dramatically increased (Figure 1B), suggesting that the majority of accumulated cholesterol in the diabetic retina is in the esterified form. A similar trend was observed for desmosterol, the final intermediate in the cholesterol biosynthesis pathway and a potent endogenous LXR agonist. In the diabetic retina, free desmosterol was decreased, and esterified desmosterol was dramatically increased (Figure 1A, B and E).

Endogenous cholesterol biosynthesis is increased in the diabetic retina. Lanosterol, an early intermediate in the cholesterol biosynthesis pathway, was elevated in both the free and total sterol pools in the diabetic cohort (Figure 1A, B). The ratio of free lanosterol to cholesterol, which is often used in plasma samples to estimate endogenous cholesterol biosynthesis, was elevated in the diabetic retina. However, while the total lanosterol concentration was increased in diabetes, the ratio of total lanosterol to cholesterol was decreased, likely due to the disproportionate accumulation of esterified cholesterol (Figure 1A, B). Interestingly, the ratio of free desmosterol to cholesterol was decreased in the diabetic cohort (Figure 1E). Given the aforementioned increase in the free lanosterol to cholesterol ratio in the diabetic retina, one would expect to observe a concomitant increase in desmosterol as a result of enhanced pathway flux. However, free desmosterol was decreased and the lanosterol to desmosterol ratio was increased (Figure 1A, B), suggesting that the diabetic retina may fundamentally alter cholesterol biosynthesis.

Impressively, DMHCA treatment corrected many of the observed shifts in the free and total sterol profiles of the diabetic cohort. Free and total desmosterol were increased with DMHCA treatment (Figure 1A, B), confirming the activity of DMHCA in the retina, which specifically inhibits desmosterol reduction leading to its accumulation. Remarkably, total cholesterol in the DMHCA treatment group was reduced by over a magnitude to baseline levels (Figure 1B, F). In addition to desmosterol, many other oxysterol species serve as endogenous LXR agonists. DMHCA treatment increased free oxysterols by >50%, including the known LXR agonist 24, 25-epoxycholesterol (Figure 1C, D). Consistent with LXR agonism, DMHCA treatment increased the transcriptional expression of the cholesterol efflux pump ABCA1 by over 100% in the diabetic retina (Figure 1G). Surprisingly, DMHCA treatment increased lanosterol and the lanosterol to cholesterol ratio, suggesting enhanced cholesterol biosynthesis (Figure 1A, B). However, the amount of free cholesterol was decreased with DMHCA treatment, meaning that any increase in cholesterol biosynthesis appears to be offset by a larger uptick in cholesterol efflux. Together, these data demonstrate the dramatic shift in the free and total sterol pools in the diabetic retina and provide strong evidence showing the beneficial effects of systemic DMHCA treatment on restoring cholesterol homeostasis in the retina.

### DMHCA rescues diabetes-induced membrane rigidity in circulating vascular reparative cells in mice and humans

In diabetes, buildup of cholesterol impedes the fluidity of the plasma membrane causing a pathologic increase in the rigidity of the cell (37, 38, 39). To test whether the observed improvements in cholesterol metabolism with DMHCA treatment could rescue this diabetic phenotype, we used an ex vivo imaging approach to quantify the effects of DMHCA on membrane fluidity of stem/progenitor cells from diabetic patients and mice. For human studies, peripheral CD34^+^ cells were collected from 19 individuals with T2D and from 19 nondiabetic control subjects (Supplemental Table 1). We chose to focus on CD34^+^ cells because membrane fluidity is especially important in this population, which require membrane flexibility to egress from the BM and a complex network of lipid rafts to transduce activation signals. Compared to the control cells (n=136), the membrane fluidity of diabetic CD34^+^ cells (n=89), as assessed by fluorescence recovery after photobleaching (FRAP), was significantly reduced (Figure 2A, B). This is consistent with the well-described dysfunction of CD34^+^ in diabetes, leading to a reduced ability to correct chronic vascular injuries such as occurs in DR. Remarkably, *ex vivo* treatment of diabetic CD34^+^ cells with DMHCA for 16-18 hours restored the fluidity of the membranes to baseline nondiabetic levels (Figure 2B). These exciting data display the potential of DMHCA to be used as a therapeutic intervention to restore the structure, and presumably function, of CD34^+^ vascular reparative cells in diabetic patients.

**Figure 2.**
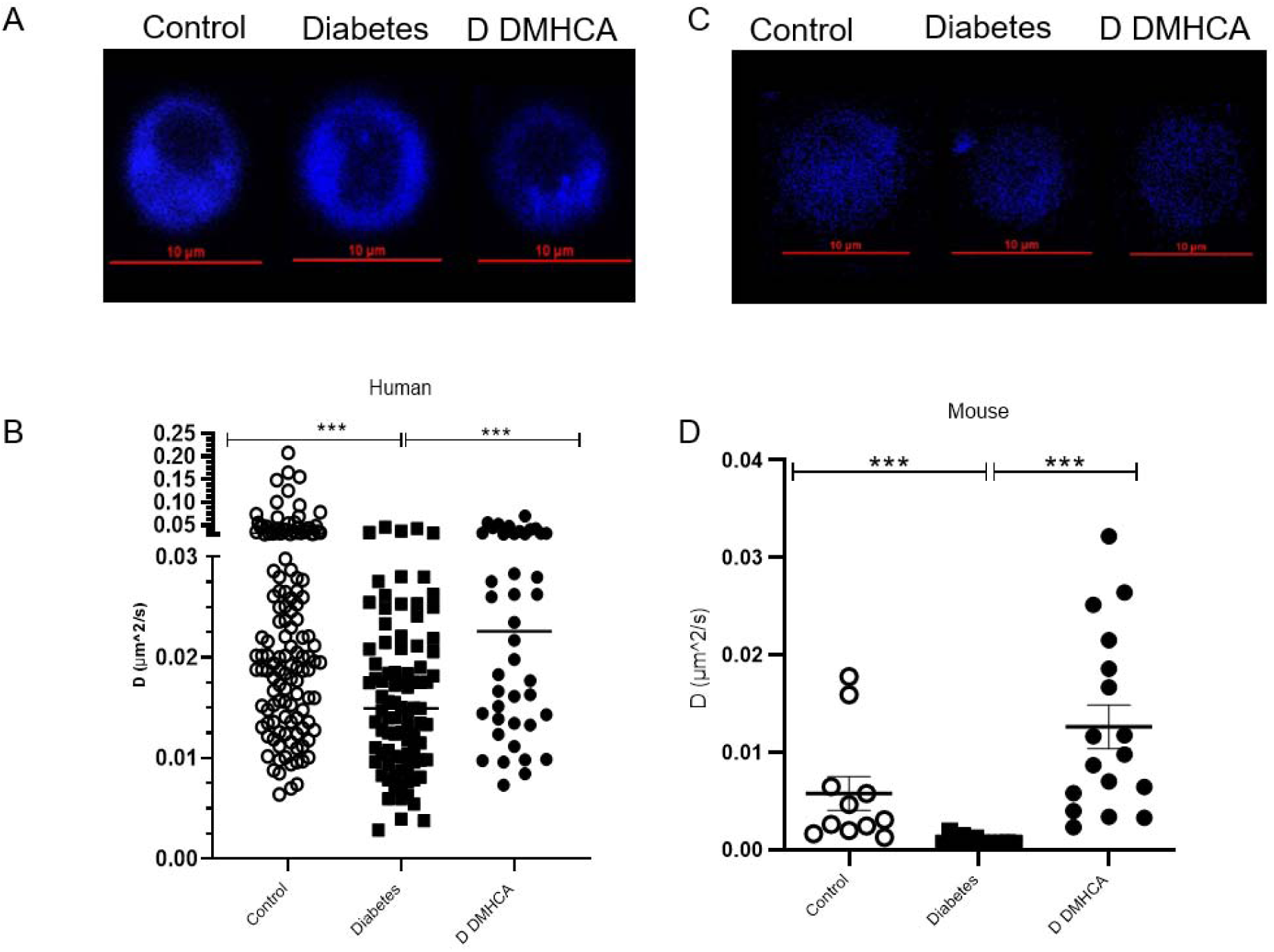
DMHCA rescues membrane fluidity in circulating vascular reparative cells in humans and mice. CD34^+^ cells were enriched from peripheral blood samples from non-diabetic (n = 19) and type 2 diabetic (n = 19) patients (**A, B**). Cells from diabetic patients were split and half received ex vivo treatment with DMHCA. Representative single-cell images of membrane staining from nondiabetic, diabetic (untreated), and DMHCA treated diabetic CD34^+^ cells (**A**). The blue color represents perylene sequesters in the membrane. Translation diffusion values from patient-derived CD34^+^ circulating vascular reparative cells. Each point on the graph represents an individual cell’s translation diffusion measurement. Control = nondiabetic cells (n=136), Diabetes = untreated diabetic cells (n=89), D DMHCA = DMHCA treated diabetic cells (n=42) (**B)**. CACs were isolated from bone marrow of nondiabetic (db/m; n =11) and diabetic (db/db; n =29) mice (**C-D**). Similar to the human studies, half of the diabetics CACs received ex vivo treatment with DMHCA. (**C**) Representative single-cell images of membrane staining from nondiabetic, diabetic (untreated), and DMHCA treated diabetic CACs. Translation diffusion values from CACs. Control = nondiabetic cells (n=11), Diabetes = untreated diabetic cells (n=12), D DMHCA = DMHCA treated diabetic cells (n=17) (**D**). Data are presented as mean ± SEM. * P < 0.05; ** P < 0.03; *** P < 0.01; analyzed using unpaired 2 tailed Mann Whitney test.

A similar approach was used to assess the *in vivo* ability of DMHCA to correct the membrane fluidity of CACs in diabetic mice. CACs were defined as CD45^+^ CD11b^-^ CD133^+^ FLK1^+^ peripheral circulating cells (Supplemental Figure 1). Compared to control cells (n=11), CACs from db/db mice (n=12) showed increased membrane rigidity, similar to what was observed in diabetic human-derived CD34^+^ cells (Figure 2C, D). After 16-18 hours of *ex vivo* DMHCA treatment, the membrane fluidity of diabetic CACs (n=17) was rescued to above baseline nondiabetic levels (Figure 2D). These data complement those observed in the human studies and demonstrate the potent ability of DMHCA to acutely correct the detrimental effects on membrane rigidity caused by diabetic dyslipidemia.

### DMHCA retards the progression of diabetic retinopathy in db/db mice

Given the remarkable ability of DMHCA to restore cholesterol homeostasis in the diabetic retina and to rescue the membrane fluidity of circulating vascular reparative cells in diabetes, we next sought to explore the functional impact of these beneficial effects on the progression of DR. Similar to humans, diabetic db/db mice develop progressive retinal pathology that shares many key features with DR including increased infiltration of pro-inflammatory leukocytes, formation of acellular capillaries, and reduced visual response (40, 41). To assess the anti-inflammatory effects of DMHCA on the diabetic retina, flow cytometry was used to quantify the relative percentages of infiltrating monocytes/macrophages. To isolate macrophages and monocytes, CD45^+^CD11b^+^ cells were gated on the macrophage marker F4-80 (Supplemental Figure 2). Macrophages were further gated on CD206 to isolate M1 CD206^-^ macrophages and M2 CD206^+^ macrophages, while monocytes were gated on CCR2 to isolate classical CCR2^-^ monocytes from nonclassical CCR2^+^ monocytes (Supplemental Figure 2). Compared to control, diabetes induced a relative increase in classical monocytes and pro-inflammatory M1 macrophages and a decrease in non-classical monocytes and reparative M2 macrophages (Figure 3A, B). Systemic DMHCA treatment rescued nearly all of these defects in the pro-inflammatory state of the diabetic retina. DMHCA reduced the relative proportion of classical monocytes to below baseline restored the proportion of non-classical monocytes to baseline, and increased the proportion of reparative M2 macrophages (Figure 3A, B).

**Figure 3.**
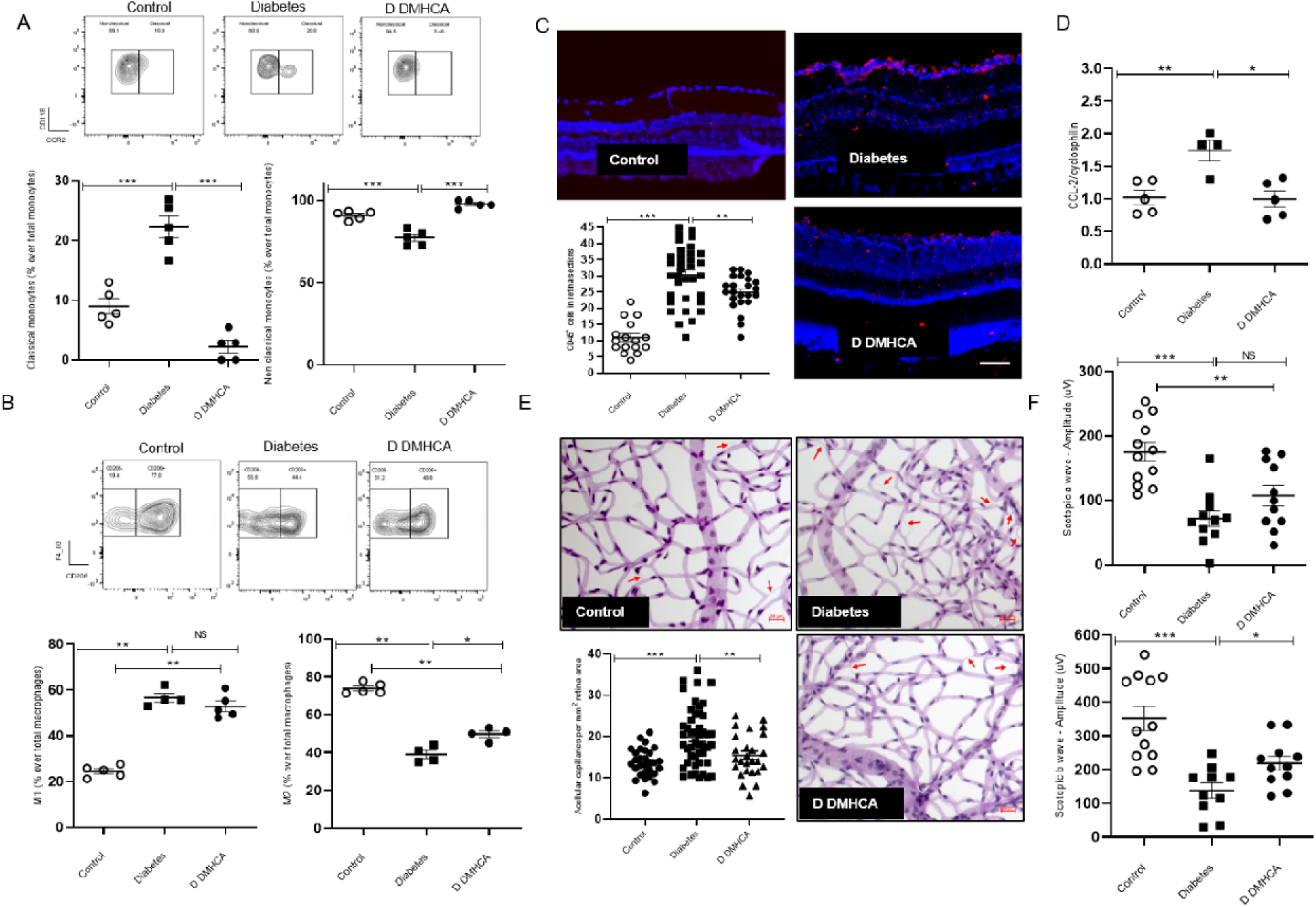
DMHCA retards the development of diabetic retinopathy in db/db mice. The presence of inflammatory and anti-inflammatory cells in the retina were determined by flow cytometry. Monocytes were defined by CD45^+^CD11b^+^Ly6G^-^ F4/80^-^ cells and classical monocytes were determined as CCR2^+^ cells and nonclassical monocytes as CCR2^-^(**A**). The macrophages were defined as CD45^+^CD11b^+^Ly6G^-^F4-80+ cells and then the CD206^-^ cells were gated to quantify M1 macrophages and the CD206^+^ to quantify M2 macrophages. (**B**). Immunofluorescence staining of retinal cross-sections for CD45^+^ cells (**C**). RT-qPCR on retinal CCL2 mRNA expression (**D**). Trypsin digested retinal flat mounts for acellular capillary quantification (**E**). Retinal visual response assessed by electroretinography. Scotopic a and b waves were quantified at an intensity of 0 db flash (3 cd/m2/s) (**F**). Data are the mean ± SEM. \**P* < 0.05; \*\**P* < 0.03 \*\*\**P* < 0.01 analyzed using unpaired 2-tailed Mann-Whitney test.

Consistent with these data, the absolute number of leukocytes in retinal cross-sections stained with CD45 were increased in untreated diabetes compared to control, and DMHCA treatment significantly reduced the number of infiltrating CD45^+^ leukocytes (Figure 3C). Furthermore, the transcript level of CCL2, a hypoxia-induced monocyte chemoattractant, was significantly increased in the untreated diabetic retina compared to control, and DMHCA restored the CCL2 transcript level to baseline (Figure 3D).

A hallmark feature of DR is microvascular dropout, which promotes retinal ischemia in diabetes. Compared to control, the number of acellular capillaries in the untreated diabetic retina was significantly increased (Figure 3E). DMHCA protected the diabetic retina from microvascular dropout, reducing the number of acellular capillaries to baseline levels (Figure 3E). These data further support the structural benefits allowed by DMHCA treatment on the diabetic retina. To assess whether these structural improvements have functional significance in DR, electroretinography was used to quantify the visual response of the retina. Compared to control, the untreated diabetic retina showed significant decreases in scotopic a- and b-waves, consistent with diabetes-induced visual dysfunction (Figure 3F). DMHCA treatment restored visual function in the diabetic animals and increased scotopic b-waves closer to baseline levels (Figure 3F). Together, these data demonstrate the beneficial effects of DMHCA treatment on the inflammatory state, vascular integrity, and visual function of the diabetic retina and suggest that DMHCA treatment has clinical potential as a treatment for DR.

### DMHCA reduces the pro-inflammatory state of the BM and increases the egression of vascular reparative cells into the peripheral circulation

In diabetes, low-level chronic inflammation alters the homeostatic balance of nearly all tissues, including the BM microenvironment (42). Myeloidosis, defined as an increase in the proportion of myeloid-derived leukocytes, is a common feature in diabetic BM and promotes the systemic inflammatory phenotype (43). Based on the observed anti-inflammatory effects of DMHCA on the diabetic retina, we asked whether the benefits afforded by systemic DMHCA treatment extended to the level of the BM microenvironment and the systemic circulation. Compared to control, cytometry bead array and ELISA analyses of BM supernatants from untreated diabetic mice displayed significant increases in the protein levels of secreted pro-inflammatory molecules, including TNF-α, IL-3, and CCL-2. Remarkably, DMHCA treatment restored the levels of BM-derived TNF-α and IL-3 to baseline and significantly reduced CCL-2 production by >50% (Figure 4A, B and C). These data are consistent with DMHCA’s ability to hamper the pro-inflammatory microenvironment of the diabetic BM.

**Figure 4.**
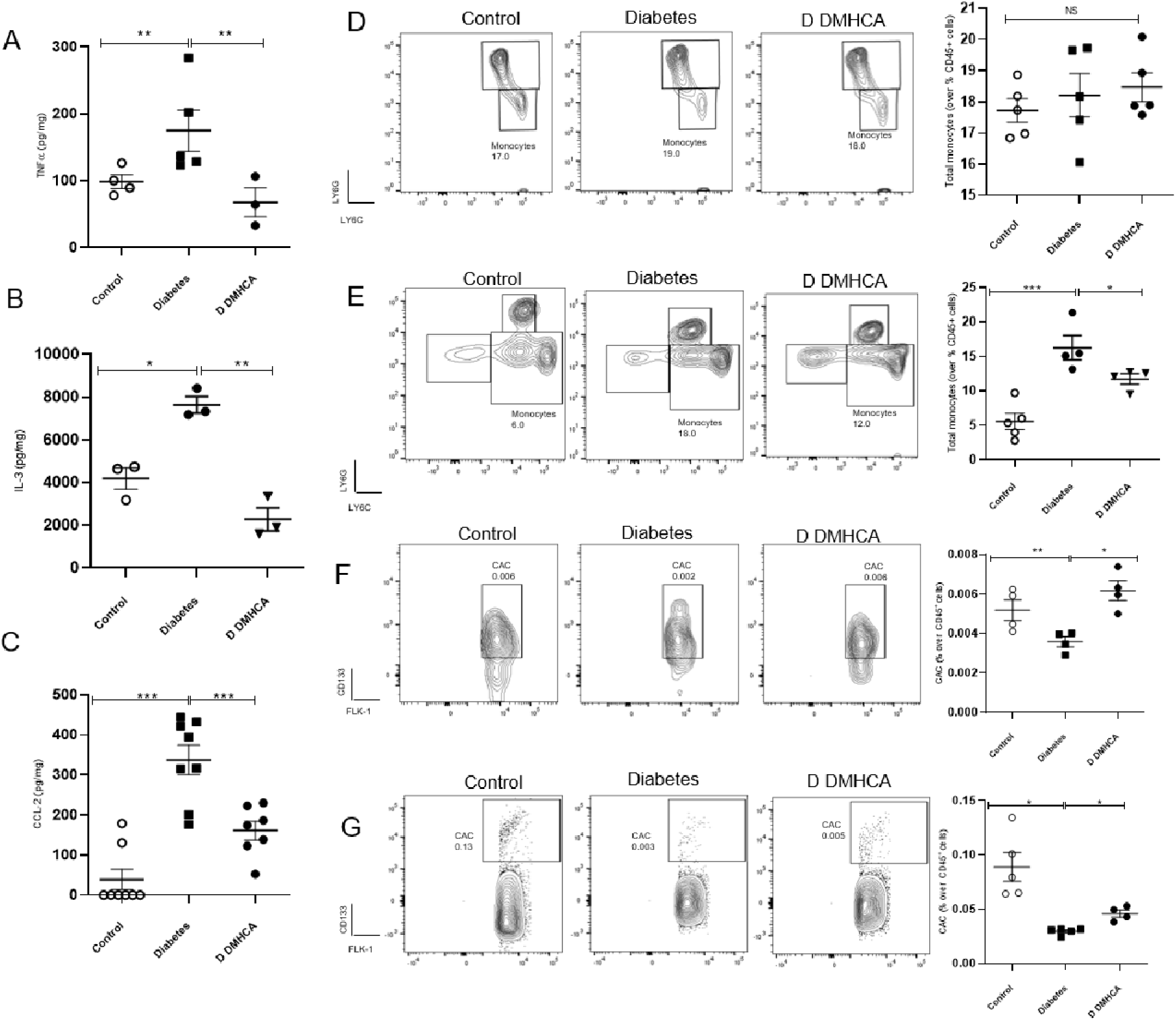
DMHCA reduces the inflammation in BM and increases the migration of vascular reparative cells into the systemic circulation. TNF-α(**A**) and CCL-2(**C**) were quantified by Cytometry beads array and lL-3 was quantified by ELISA(**B**). Monocytes from bone marrow (**D**) and peripheral blood (**E**) were quantified by flow cytometry and defined as CD45^+^CD11b^+^Ly6G^-^LY6C^+^ cells. Flow cytometry analysis of levels of circulating angiogenic cells (CAC) were determined as CD45^+^CV11b^-^CD133^+^FLK1^+^ cells of bone marrow cells(**F**) and peripheral blood (**G**). Data are the mean ± SEM. \**P* < 0.05; \*\**P* < 0.03 \*\*\**P* < 0.01 analyzed using unpaired 2-tailed Mann-Whitney test.

Next, flow cytometry was used to profile specific populations of leukocytes in the BM and peripheral circulation (Supplemental Figure 3). Compared to control, diabetes induced a significant increase in the number of circulating monocytes but surprising, had no effect on the total monocyte population of the BM (Figure 4D). DMHCA treatment significantly reduced the number of circulating monocytes in diabetic mice (Figure 4E). We next focused on vascular reparative cells in the BM and circulation. As previously reported, diabetes induced a significant decrease in the total proportion of CACs in the BM and peripheral circulation (Figure 4F, G). Compared to untreated diabetic mice, DMHCA significantly increased the number of CACs in the BM and peripheral circulation (Figure 4F, G). These findings complement the previous data showing enhanced membrane fluidity in DMHCA-treated CACs and suggests that the improvements in membrane fluidity may account for the observed increase in CAC egression into the peripheral circulation. Taken together, these data demonstrate that systemic DMHCA treatment has the additional benefit of preventing diabetes-induced myeloidosis and enhancing the egression of vascular reparative cells into the peripheral circulation.

### DMHCA acts at the level of HS/PC to correct diabetic myeloidosis

To better understand the mechanism by which DMHCA normalizes the composition of circulating peripheral leukocytes, we next focused on the hematopoietic stem cell (HSC) compartment. During hematopoiesis, differentiation signals instruct HSCs to favor specific lineages, and a homeostatic balance of these signals is necessary to maintain equilibrium of circulating cells. In certain pathologic states, this balance becomes uneven leading to the accumulation of a particular lineage, such as in diabetic myeloidosis. To explore the HSC compartment, we gated on lineage^-^, SCA1^+^, c-KIT^+^ (LSK) BM cells (Supplemental Figure 4). CD34^+^FLT3^+^ multi-potent progenitors (MPPs) accounted for the largest majority of LSK cells, followed by CD34^+^FLT3^-^ short term HSCs (ST-HSCs) and finally CD34^-^FLT3^-^ long term HSCs (LT-HSCs). Compared to controls, untreated diabetic LSK cells displayed decreased ST-HSCs and LT-HSCs and increased MPPs (Figure 5A, B and D). In the treated diabetic cohort, DMHCA significantly reduced the proportion of MPPs back to baseline and increased LT-HSCs (Figure 5A, C and D). These data support the notion that DMHCA is sufficient to restore the HSC compartment in diabetes towards a nondiabetic state.

**Figure 5.**
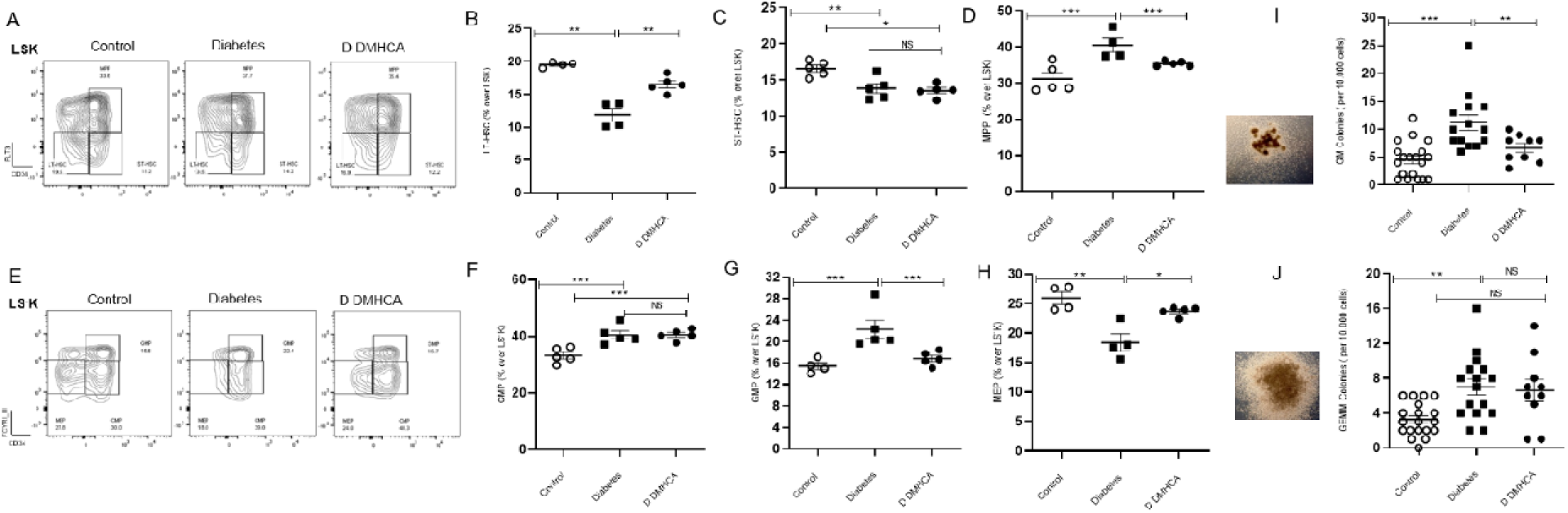
DMHCA corrects hematopoietic stem and progenitor dysfunction in db/db mice. BM HSC (**A-D**) and HPC (**E-H**) was assessed by flow cytometry. HSCs were defined as Lin^-^ Sca1^+^ c-Kit^+^ (LSK) BM cells (**A**). LT-HSCs were defined as CD34^-^FLT3(**B**) ^-^, ST-HSCs as CD34^+^FLT3^-^(**C**), and MPPs as CD34^+^FLT3^+^(**D**). HPCs were defined as Lin^-^ Sca1^-^ c-Kit^+^ BM cells (**E**). CMPs were defined as CD32/CD16^-^ CD34^+^(**F**), GMPs as CD32/CD16^+^ CD34^+^(**G**), and MEPs as CD32/CD16^-^ CD34^-^(**H**). Colony forming unit (CFU) assays for ex vivo differentiation of cultured HPCs**(I-J)**. Granulocyte, macrophage (GM; bottom image) CFUs **(I)** and granulocyte, erythroid, macrophage, megakaryocyte (GEMM; bottom image) CFUs (**J**). Data are presented as mean ± SEM. * P < 0.05; ** P < 0.03; *** P < 0.01; analyzed using unpaired 2 tailed Mann Whitney test.

Lineage-committed progenitor populations (lineage^-^sca1^-^ckit^+^) were examined next to determine the effect of DMHCA on hematopoietic lineage flux (Figure 5E). CD32/CD16^-^CD34^+^ common myeloid progenitors (CMPs) accounted for the largest percentage of cells, followed by CD32/CD16^+^CD34^+^ granulocyte myeloid progenitors (GMPs) and finally CD32/CD16^-^CD34^-^ megakaryocyte-erythrocyte progenitors (MEPs). Untreated diabetic mice showed an increase in CMPs and GMPs and a decrease in MEPs, consistent with diabetic myeloidosis (Figure 5F, G and H). Again, DMHCA treatment corrected many of the defects observed in the lineage-committed progenitor populations in diabetes. DMHCA significantly reduced the number of GMPs, suggesting a decrease in the production of neutrophils and monocytes (Figure 5G). This is consistent with our previous observations demonstrating reduced circulating monocytes with DMHCA treatment and suggests that the mechanism relates, at least partially, to the ability of DMHCA to correct diabetic myeloidosis. In addition, DMHCA treatment significantly increased the proportion of MEPs to baseline levels, suggesting an increase in the production of erythrocytes and megakaryocytes (Figure 5H).

Finally, we assessed the differentiation ability of BM-derived stem and progenitor cells using an *ex vivo* culture assay. BM from control, diabetic, and DMHCA-treated mice were enriched for progenitor markers and grown in culture for 12 days, after which the number of colony forming units (CFU) was counted. In BM from untreated diabetic mice, there were significantly more granulocyte, erythroid, macrophage, megakaryocyte (GEMM) CFUs and granulocyte, macrophage (GM) CFUs compared to control, again consistent with diabetic myeloidosis (Figure 5I). In diabetic mice treated with DMHCA, the number of GM-CFUs was reduced to baseline levels while no effect was observed on GEMM-CFUs (Figure 5J). Together, these data suggest that DMHCA acts at the level of hematopoietic stem and progenitor cells to fundamentally shift diabetic hematopoiesis towards a more normal nondiabetic state. Remarkably, this effect is sufficient to suppress diabetic myeloidosis.

### Single-cell analysis of diabetic HSCs in DMHCA treated mice

To gain mechanistic insight into DMHCA’s ability to influence diabetic HSCs and progenitors, single-cell RNA-seq was performed on LSK sorted BM cells. 5,103 cells were recovered from the untreated diabetic group and 5,152 cells from the DMHCA treated group, for a total of 10,255 HSCs. Unsupervised clustering was performed on the normalized, batch corrected, and cell-cycle gene regressed data and revealed 13 distinct clusters (Figure 6A). The largest population of cells, accounting for 27% of the total combined sample, was multi-potent stem cells (Figure 6E). These cells were identified by high expression of the stem cell markers CTLA2A, HLF, and CD34 and the absence of lineage specific gene expression (Figure 6B). Gene expression patterns of all major lineages were represented in the single-cell analysis, albeit at varying levels. These include dendritic cell progenitors (CD74, H2-AA, and H2-EB1 high), erythrocyte progenitors (HBB-BT high), lymphoid progenitors (DNTT high), monocyte progenitors (IRF8 and LY86 high), megakaryocyte/basophil progenitors (VWF and GATA2 high), neutrophil progenitors (MPO and CTSG high), and pre-B and T cell progenitors (EBF1/CD19 high and TRBC1 high, respectively) (Figure 6B, C).

**Figure 6.**
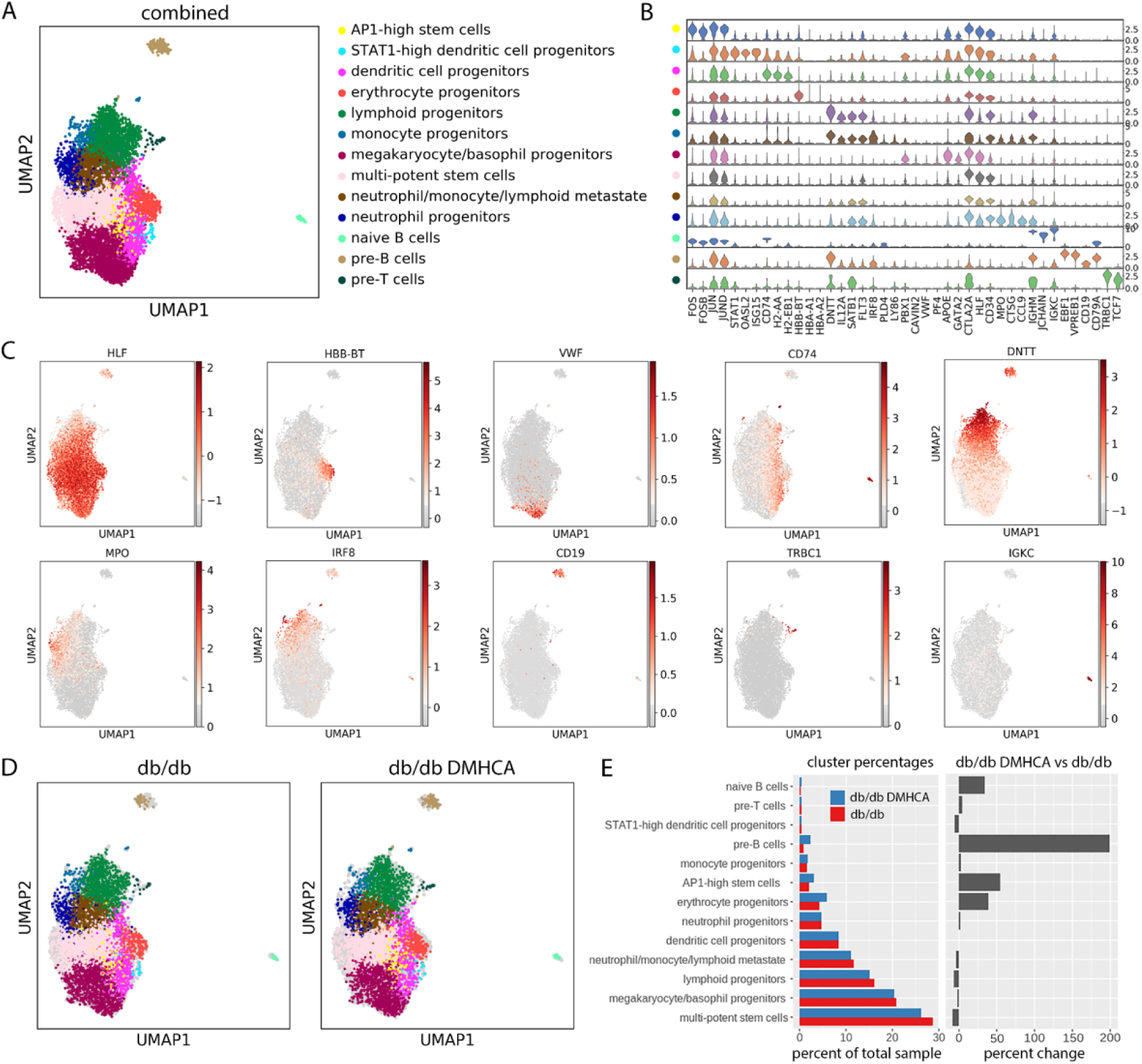
Single-cell RNA-seq analysis of untreated and DMHCA treated diabetic HSCs. UMAP representation of scRNA-seq from LSK sorted HSCs reveals 13 distinct clusters (**A**). Violin plots of lineage-specific gene expression across all 13 clusters (**B**). Spatial representation of lineage-specific gene expression (**C**). UMAP cluster representations of untreated (db/db) and DMHCA treated (db/db DMHCA) diabetic HSCs (**D**). Cluster proportions of all 13 cell populations in untreated and DMHCA treated samples (**E**).

While the cluster distributions appear similar between the two groups (Figure 6D), differences are noted in the proportions of individual clusters (Figure 6E). In the DMHCA treatment group, multi-potent stem cells decreased by 8.8% (Figure 6E). This is consistent with our previous finding that DMHCA reduces the pathologic increase of MPPs in diabetic BM. Additionally, DMHCA increased the relative proportion of erythrocyte progenitors by 39% (Figure 6E). This is consistent with our previous observation of increased MEPs in DMHCA-treated BM. Interestingly, a novel population of stem cells was identified, which we refer to as AP1-high stem cells. This relatively small population, accounting for roughly 3% of the total sample, was identified as expressing high levels of the activator protein 1 (AP1) complex, including FOS, FOSB, JUN, JUNB, and JUND, and largely lacked expression of lineage specific genes (Figure 6B). In the DMHCA treatment group, this AP1-high stem cell population was increased by 54% (Figure 6E). Moreover, DMHCA treatment unexpectedly increased the relative proportion of both pre-B and naïve B cells (Figure 6E). Lastly, of note is the lack of change observed in the neutrophil and monocyte progenitor populations (Figure 6E). This suggests that DMHCA’s ability to influence these cells and correct diabetic myeloidosis may occur at a later stage in hematopoiesis.

### DMHCA treatment increases expression of immediate early response genes

To probe the transcriptional pathways responsible for DMHCA’s effect on diabetic HSCs, we performed differential gene expression (DGE) analysis on the total DMHCA treated group compared to the untreated control. Using an FDR cutoff of 0.01 and accounting for technical covariates, we identified 1,048 differentially expressed genes (Figure 7A). The majority of DGEs were increased with DMHCA and included, among others, genes associated with immediate early response (FOS, FOSB, FOSL2, JUN, JUNB, JUND, ATF4, EGR1, EGR3, MYC, IER3, MCL1), LXR activation (APOE, FASN, PTGES3, HNRNPAB, SLC3A2, ETF1, RANBP1, PRDX2), and lineage-specification (HBB-BT, CD19) (Figure 7B). Pathway enrichment analysis identified the three most highly upregulated pathways as EIF2 signaling, elF4/p70S6K signaling, and mTOR signaling (Figure 7C). Notably, the NRF2 oxidative stress response, hypoxia signaling, and oxidative phosphorylation were also upregulated (Figure 7C).

**Figure 7.**
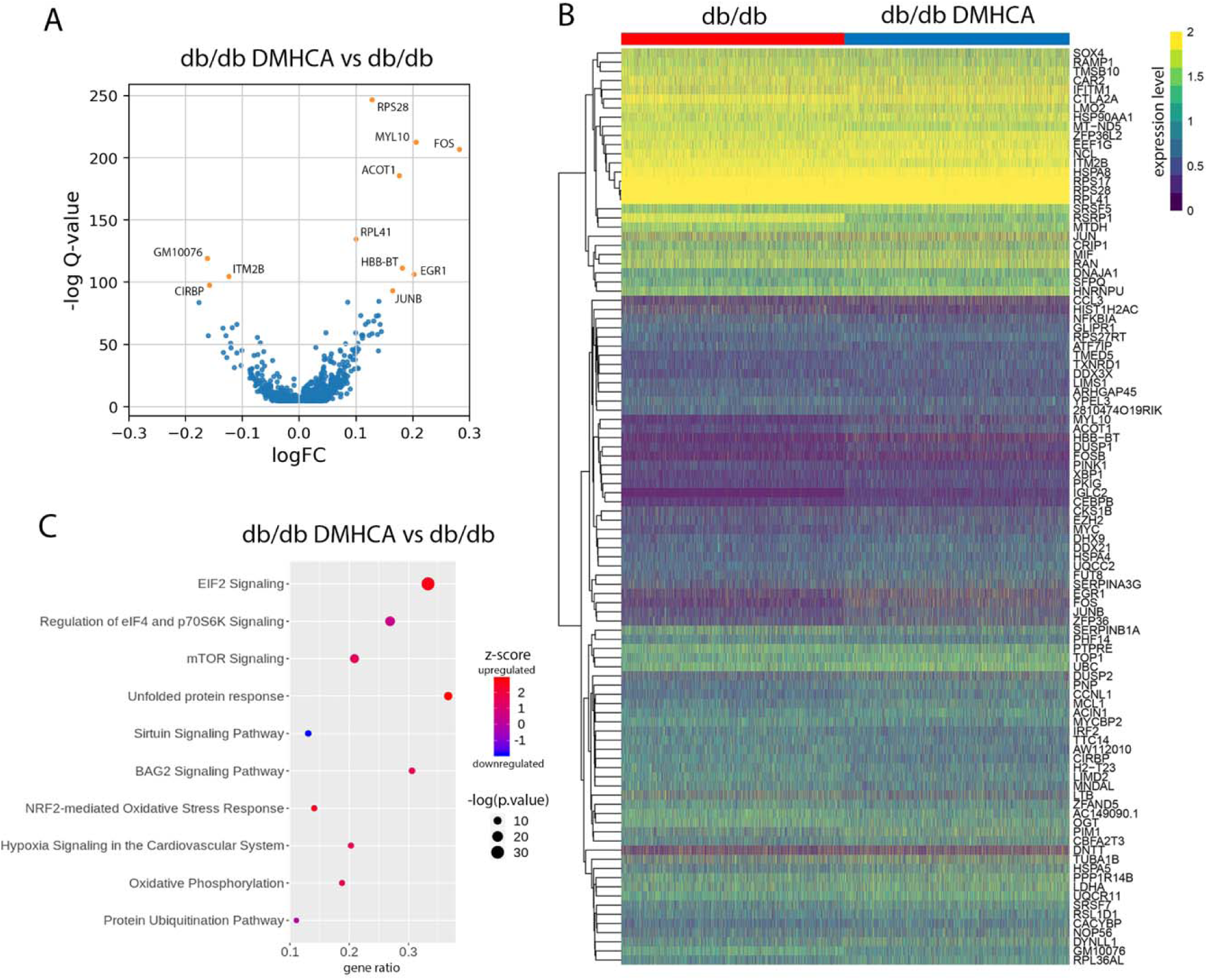
DMHCA treatment increases expression of immediate early response genes. Differentially expressed genes in DMHCA treated diabetic HSCs (**A**). Heatmap of the top 100 log(fold change) genes from the DGE analysis(**B**). Top 10 pathways from IPA pathway enrichment analysis(**C**).

### DMHCA treatment enhances flux down the erythrocyte progenitor lineage

Similar to most differentiation processes, hematopoiesis proceeds across a spectrum of gene expression changes rather than in discrete discernible steps. Thus, single-cell profiles of HSCs represent individual snapshots in time, where each cell falls somewhere along the differentiation spectrum. Pseudotime trajectory analysis relies on the identification of specific gene expression patterns within the dataset to map the trajectory of cells along specific lineages. Using this technique, we identified two main differentiation pathways, one leading to megakaryocytes, erythrocytes, and dendritic cells, and another to peripheral circulating leukocytes (Figure 8A). Partition-based graph abstraction (PAGA) analysis uses a similar approach to physically map cells along a spectrum of gene expression changes and provides enhanced resolution. PAGA analysis of the combined dataset revealed a similarly distinct split between megakaryocytes/basophils and erythrocytes and peripheral leukocytes, including lymphoid, monocyte, neutrophil, pre-B, and pre-T progenitors (Figure 8B). Differentiation along the peripheral leukocyte division mostly precedes through a transition state characterized by low-level expression of several lineage-specific genes, which we refer to as a neutrophil/monocyte/lymphoid metastate (Figure 8C). This metastate then trifurcates into neutrophil, lymphoid, and pre-T cell progenitors (Figure 8C). Finally, monocyte progenitors derive from the neutrophil progenitor population, while pre-B cells derive from lymphoid progenitors (Figure 8C).

**Figure 8.**
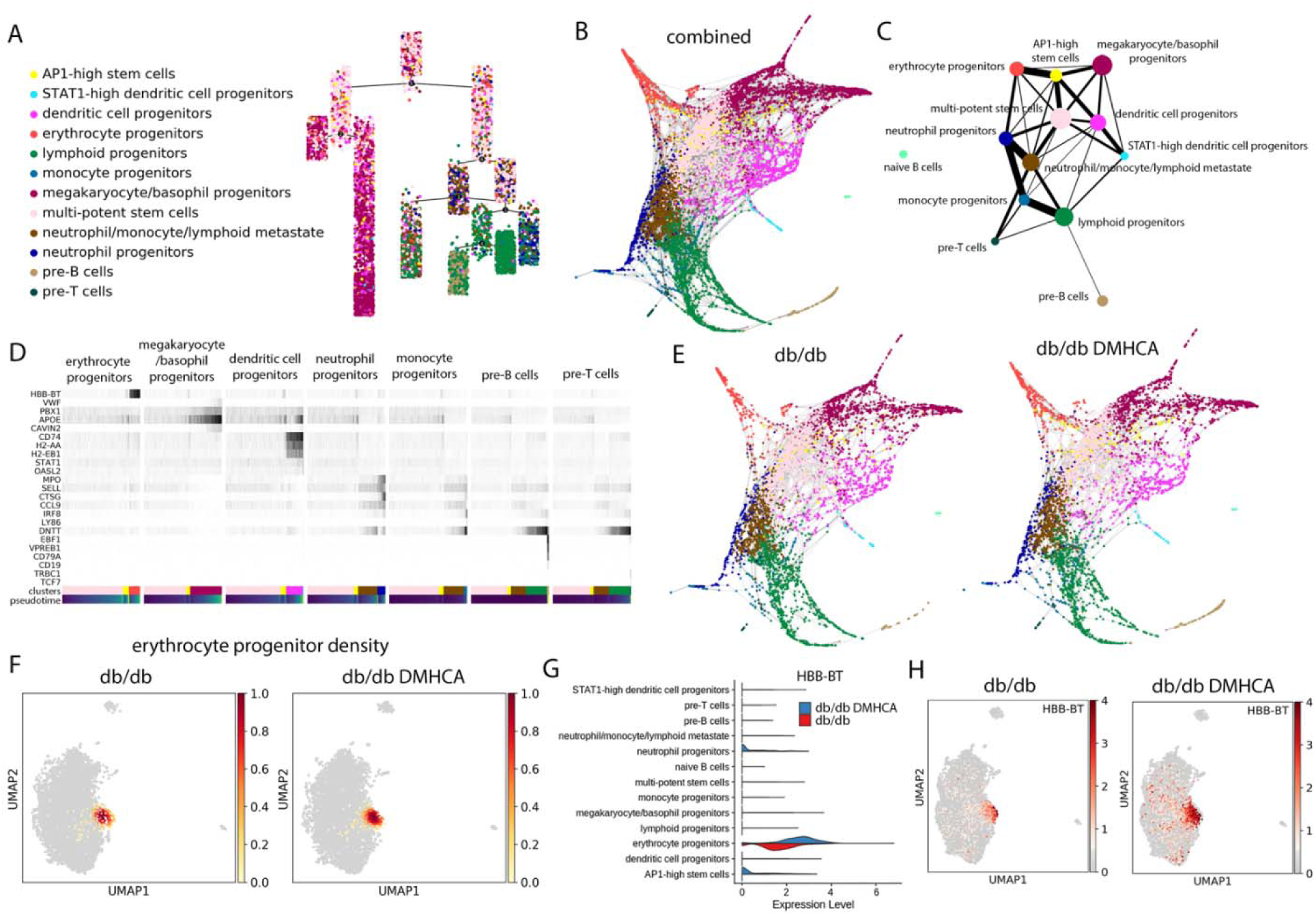
Trajectory analysis of HSC differentiation reveals increased erythrocyte progenitor flux with DMHCA treatment. Pseudotime trajectory analysis identifies early partitioning of two distinct HSC differentiation pathways (**A**). ForceAtlas2 embedding of PAGA analysis (**B**). Ball-and-stick representation of PAGA analysis. Circle size represents number of cells and line thickness represents connectivity between two groups of cells (**C**). Gene expression changes of lineage-specific genes along pseudotime differentiation of seven lineages (**D**). ForceAtlas2 embedding of PAGA analysis separated by sample (**E**). Compositional analysis showing density graphs of erythrocyte progenitors in untreated and DMHCA treated diabetic HSCs (**F**). Violin plots comparing HBB-BT gene expression across samples(**G**). Spatial representation of HBB-BT gene expression across samples (**H**).

Interestingly, our analysis suggests that differentiation along the dendritic lineage is more closely related to the megakaryocyte/basophil and erythrocyte family and may actually represent a distinct division arising directly from the HSC compartment (Figure 8C). In addition, we further analyzed the novel AP1-high stem cell population to better understand which lineages these cells contribute to. PAGA analysis revealed that AP1-high stem cells are derived directly from HSCs and predominantly give rise to erythrocyte progenitors and, to a lesser degree, megakaryocyte/basophil and dendritic cell progenitors (Figure 8C). Differentiation along the aforementioned lineages occurs through a gradual increase in lineage-specific genes (Figure 8D).

Lastly, we separated the PAGA analysis into the two discrete sample groups and found that DMHCA enhanced flux down the erythrocyte progenitor lineage (Figure 8E). Compositional analysis comparing cellular densities between conditions confirmed the increase in erythrocyte progenitor density in DMHCA-treated BM (Figure 8F). Impressively, DMHCA not only expanded the proportion of erythrocyte progenitors but also increased the expression of the hemoglobin beta adult t chain (HBB-BT) (Figure 8G, H). Together, these data suggest that, at the earliest stage of hematopoiesis, DMHCA treatment enhances the overall production and robustness of erythrocyte progenitors.

### DMHCA enhances signaling in the AP1-high stem cell and erythrocyte progenitor populations

We next focused on the AP1-high stem cell and erythrocyte progenitor populations to better understand how DMHCA influences these clusters. Compositional analysis confirmed an increase in the density of the AP1-high stem cell population with DMHCA treatment (Figure 9A). Based upon our trajectory analysis, which identified the AP1-high stem cell population as precursors to erythrocyte progenitors, the increase in AP1-high stem cells is consistent with the observed enhancement in erythrocyte flux. We next performed a differential gene expression analysis comparing the AP1-high stem cell populations in untreated and DMHCA treated mice. Owing to the relatively small number of cells in this population (261 cells total), only 9 genes were found to be differentially expressed and included, among others, the AP-1 genes FOS and FOSB and the Krüppel-like family of transcription factors (KLFs) KLF6 and KLF2 (Figure 9B). Pathway enrichment analysis focusing on intracellular and secondary messenger signaling identified the ERK and MAPK pathways as the most enriched, followed by glucocorticoid and JAK/STAT signaling (Figure 9C).

**Figure 9.**
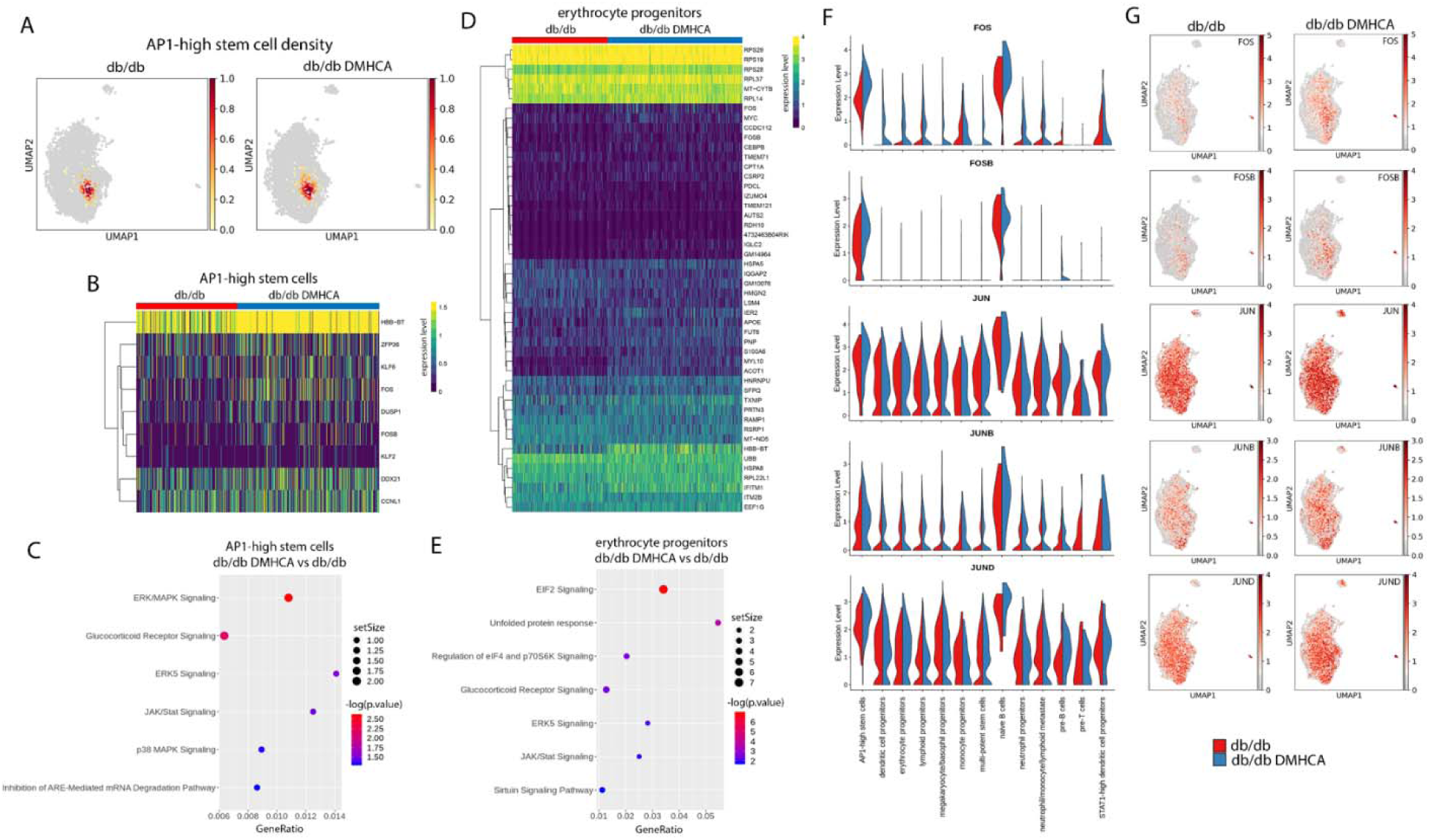
DMHCA induces subpopulation gene expression changes and enhances AP-1 signaling. (**A**) Compositional analysis showing density graphs of AP-1 high stem cells in untreated and DMHCA treated diabetic HSCs (**B**). Heatmap of differentially expressed genes in AP1-high stem cells from untreated and DMHCA treated HSCs (**C**). Significantly enriched secondary/intracellular signaling pathways in DMHCA treated AP1-high stem cells from IPA pathway enrichment analysis (**D**). Heatmap of differentially expressed genes in erythrocyte progenitors from untreated and DMHCA treated HSCs (**E**). Significantly enriched secondary/intracellular signaling pathways in DMHCA treated erythrocyte progenitors from IPA pathway enrichment analysis (**F**). Violin plots comparing expression of AP-1 complex genes across samples and clusters (**G**). Spatial representation of AP-1 complex gene expression across samples.

Next, we performed a differential gene expression analysis comparing the erythrocyte progenitor populations in untreated and DMHCA treated mice. Across 517 cells, 48 differentially expressed genes were identified (Figure 9D). Of note, these included genes involved in the immediate early response (FOS, FOSB, MYC, and IER2), hemoglobin synthesis (HBB-BT), and ribosome synthesis (RPS29, RPS19, RPS28, RPL37, and RPL14) (Figure 9D). Using these 48 differentially expressed genes, we performed pathway enrichment focusing on intracellular and secondary messenger signaling. Similar to the AP1-high stem cell population, glucocorticoid, ERK5 and JAK/STAT pathways were significantly enriched (Figure 9E). However, unlike the AP1-high population, the largest enrichment in erythrocyte progenitors was related to translation initiation (EIF2 and EIF4/p70S6K signaling) (Figure 9E).

Lastly, we examined the distribution of AP-1 genes across the 13 clusters of the untreated and DMHCA-treated diabetic mice. FOS, which encodes the c-Fos protein, is predominantly expressed in the AP1-high stem cell and naïve B cell populations (Figure 9F, G). DMHCA increased FOS expression in nearly all clusters, including most strongly the AP1-high stem cell and naïve B cell populations (Figure 9F, G). FOSB expression is highly specific to the AP1-high stem cell and naïve B cell populations, and DMHCA increased FOSB in these two clusters (Figure 9F, G). JUNB is expressed to varying degrees in all clusters but is predominantly found in the AP1-high stem cell and naïve B cell populations (Figure 9F, G). DMHCA treatment increased JUNB expression in all clusters, especially the AP1-high stem cell and naïve B cell populations (Figure 9F, G). Lastly, JUN (c-Jun) and JUND are highly expressed in all clusters, and DMHCA caused a pan-increase in their transcriptional expression (Figure 9F, G).

## Discussion

In this study, we use a multi-system approach to demonstrate the remarkable efficacy of the novel LXR agonist, DMHCA, on correcting diabetic dysfunction related to the retina-BM axis. In aged diabetic db/db mice, DMHCA treatment restores retinal cholesterol homeostasis, retards the development of DR, hampers chronic systemic inflammation, and corrects BM dysfunction. In circulating vascular reparative cells from diabetic patients and db/db mice, DMHCA rejuvenates membrane fluidity and promotes BM egression. Given the selective nature of DMHCA’s mechanism of LXR agonism, which promotes cholesterol efflux and hampers systemic inflammation without inducing hypertriglyceridemia, the results herein provide compelling evidence that DMHCA treatment has the potential to provide multi-system protection in diabetic patients.

Our studies using highly sensitive quantitative LC-MS to measure free and total sterols in the diabetic retina provide an unprecedented view of the dysfunctional cholesterol homeostasis in diabetes. From these results, many novel conclusions are supported. First, in the diabetic retina, cholesterol buildup is immense and results from the accumulation of esterified cholesterol. This important finding is supported by the observation that free cholesterol is actually decreased in diabetes, while total cholesterol (total = free + esterified) is profoundly increased by 1.5 magnitudes. Second, desmosterol, a potent endogenous LXR agonist, follows a similar pattern to cholesterol in the diabetic retina, where free desmosterol is decreased and esterified desmosterol is dramatically increased. Esterification of desmosterol involves the addition of a long fatty acid tail, chemically altering the composition of desmosterol and likely reducing its ability to activate LXR, which requires side chain hydrogen bond acceptors for potent activation (10). This finding potentially explains the previously observed decreases in LXR activity and transcript levels in the diabetic retina (18), which are autoregulated by LXRα activation (36). Third, diabetes alters the lanosterol to desmosterol ratio and surprisingly decreases free desmosterol in the face of enhanced cholesterol biosynthesis. This finding is unexpected given the increased flux in the cholesterol biosynthesis pathway, which would be presumed to increase free desmosterol, and suggests that diabetes may fundamentally alter retinal cholesterol biosynthesis. In classical cholesterol biosynthesis, two separate pathways are responsible for cholesterol synthesis, the Bloch pathway and the Kandutsch-Russell pathway. The Bloch pathway involves the production of the intermediate desmosterol as the final precursor to cholesterol, while the alternative Kandutsch-Russell pathway uses 7-dehydrocholesterol as its final precursor. Different tissues utilize differing ratios of the two pathways during cholesterol biosynthesis, suggesting that there are cell-specific mechanisms responsible for the homeostatic balance of the two. Given the decreased lanosterol to desmosterol ratio in the diabetic retina, this finding suggests that diabetes may fundamentally shift retinal cholesterol biosynthesis to a more Kandutsch-Russell predominant pathway, thereby accounting for the decreased production of active free desmosterol. Remarkably, systemic DMHCA treatment is sufficient to correct nearly all of these pathologic alterations in cholesterol homeostasis in the diabetic retina.

To test whether the lipidomic benefits afforded by DMHCA treatment are sufficient to provide structural and functional benefit in diabetes, we assessed retinal function and membrane fluidity in circulating vascular reparative cells derived from diabetic patients and mice. We found that DMHCA-treatment corrected several DR endpoints in db/db mice, including reduced vascular dropout and enhanced visual function. Remarkably, in diabetic human derived CD34^+^ cells, acute *ex vivo* treatment with DMHCA was sufficient to restore membrane fluidity, suggesting that DMHCA has potential as a novel therapeutic approach to correcting diabetes-induced dysfunction of circulating vascular reparative cells. Interestingly, DMHCA treatment also reduced leukocyte trafficking to the diabetic retina, providing compelling evidence of DMHCA’s anti-inflammatory effects in diabetes.

To test whether DMHCA’s anti-inflammatory effects extend to the level of the peripheral circulation and BM, we assessed these compartments for inflammatory markers and quantified the proportions of specific cell populations. DMHCA treatment significantly reduced the expression of several important pro-inflammatory proteins in the BM, including TNF-α, IL-3, and CCL-2. Moreover, DMHCA treatment significantly reduced circulating monocytes and increased the proportion of vascular reparative cells in the BM and circulation. Vascular reparative cells are particularly sensitive to alterations in membrane fluidity, as they require flexibility to egress into the circulation and complex intercellular signaling networks to home to areas of injury. These findings on the increased production and egression of CACs complement our membrane fluidity results and suggest that the improvements in membrane fluidity have functional benefit.

Given the beneficial effects of DMHCA on correcting the homeostatic balance of BM-derived cells, we next explored the HSC and BM progenitor compartments to test whether DMHCA treatment directly influences hematopoiesis. In LSK cells, DMHCA treatment was sufficient to restore the diabetes-induced increase in MPPs and decrease in LT-HSCs. Moreover, DMHCA reduced granulocyte myeloid progenitor hyperplasia and increased the production of megakaryocyte/erythrocyte progenitors. Lastly, DMHCA treatment was sufficient to lower *ex vivo* granulocyte/macrophage CFUs to baseline levels. Taken together, these exciting findings suggest that DMHCA’s benefits extend to the hematopoietic stem cell compartment and that DMHCA treatment is sufficient to correct diabetic myeloidosis.

Lastly, to better understand the transcriptional mechanisms responsible for DMHCA’s beneficial effects on diabetic hematopoiesis, we performed single-cell RNA-seq (scRNA-seq) on untreated and DMHCA treated LSK cells. The findings from our scRNA-seq analysis have important implications for our understanding of early fate decisions in HSCs. One striking finding in our analysis is that even the most primitive hematopoietic cells appear pre-programmed to enter specific lineages. This is supported by the 13 cell populations identified by unsupervised clustering, which account for nearly all of the main lineages derived from BM. Another exciting finding from our scRNA-seq analysis is that dendritic cell differentiation appears to be more nuanced than previously suggested. A recent single-cell report using c-Kit^+^ murine BM found that dendritic and monocytes lineages split late in differentiation (44). Our trajectory analysis, which uses a more primitive population of cells and thus provides a higher resolution of early fate decisions, suggests that the dendritic lineage is more closely related to the megakaryocyte/basophil and erythrocyte family and may actually represent a distinct division arising directly from the HSC compartment. Interestingly, in another recent scRNA-seq report on murine LSK sorted HSCs, the authors did not mention the dendritic cell lineage (45). However, inspection of their publicly available dataset for expression of early dendritic cell markers (CD74, H2-AA, H2-EB1) - as defined by Tusi et al (44) - demonstrated close association of these genes with the HSC population (Supplemental Figure 5), thus supporting our results herein. This finding on the dendritic cell lineage may have important implications as it suggests that the development of an entire arm of the immune system may be fundamentally different from all other immune cells, which in our analysis, derive from a shared hematopoietic branch. Lastly, our analysis identifies a novel and unique stem cell population, which we refer to as AP1-high stem cells. This relatively small population of cells express disproportionately high levels of AP-1 complex genes and largely lack lineage specification. Trajectory analysis suggests that AP-1 high stem cells predominantly give rise to erythrocyte progenitors and to a lesser degree, megakaryocyte/basophil and dendritic cell progenitors. Thus, this cluster may represent a novel HSC population that is an intermediate to the aforementioned lineages.

The findings from our scRNA-seq analysis also help to elucidate DMHCA’s mechanism of action in the HSC compartment. The main findings from our DMHCA treated sample were a decrease in the multi-potent stem cell population and an increase in the AP1-high stem cell and erythrocyte progenitor populations. Both of these findings are supported in our BM HSC studies, which found that DMHCA suppressed diabetes-induced hyperplasia of multipotent progenitors and increased production of erythrocyte/megakaryocyte progenitors. Based upon our trajectory analysis, which identified the AP1-high stem cell population as precursors to erythrocyte progenitors, the increase in AP1-high stem cells is consistent with the observed enhancement in erythrocyte progenitor flux. DGE analysis demonstrated that DMHCA increased the expression of several LXR target genes (46), confirming that DMHCA fact directly modulates these primitive cells. Moreover, we observed a striking increase in immediate early response genes. These gene targets, such as FOS, EGR1, and MYC, are pleiotropic factors involved in many cell processes including differentiation. They are termed immediate genes because they are rapidly induced in response to inter- and intra-cellular signaling (47). Many of these genes have well-characterized roles in hematopoiesis. For example, FOS expression is known to limit HSC hyperplasia (48), whereas FOS depletion results in a >90% reduction of clonogenic B-cell precursors (49). Both of these observation are consistent with our results herein. Given our previous findings on improved membrane fluidity with DMHCA treatment, the increase in immediate early gene expression in HSCs is not surprising as the enhanced formation of membrane microdomains amplifies the transduction of intercellular signaling. The most highly-enriched pathways in DMHCA treated HSCs were related to ribosome synthesis/translation initiation (EIF2 and EIF4/p70S6K signaling) and mTOR signaling. Moreover, pathway enrichment analysis indicated upregulation of the hypoxia and Nrf-2 pathways. At normal steady-state, the BM microenvironment is hypoxic, and cellular stress stimulates increase BM pO_2_ (50). Nrf2 is the master regulator of several cellular antioxidant pathways. As Nrf2 expression is decreased in diabetic BM cells (51), DMHCA induced upregulation of Nrf-2 may function as a pro-survival response. Together, these findings suggest that DMHCA treatment normalizes the BM microenvironment.

Lastly, we found key genes and intracellular pathways that were differentially regulated in the treated versus untreated AP1-high stem cell and erythrocyte progenitor populations. DMHCA treated AP1-high stem cells showed increased AP-1 complex genes as well as increased expression of KLF2 and 6. Interestingly, homozygous knockout of either KLF2 or KLF6 is embryonic lethal and required for normal erythrocyte development (52, 53). Pathway enrichment analysis found significantly enriched ERK and MAPK pathways, consistent with immediate early gene expression (47). DMHCA treated erythrocyte progenitors showed induction of ribosomal and immediate response genes. Of note is the increase in MYC expression, which has recently been shown to induce proliferation of erythroid progenitor cells in an ex vivo method used to produce large quantities of red blood cells (54). DGE analysis also showed significantly increased hemoglobin gene expression, suggesting that DMHCA not only increases erythrocyte progenitor flux but also stimulates the production of more robust erythrocyte progenitors. Pathway analysis showed significantly enriched ERK signaling as well as translation initiation pathways, consistent with immediate gene response. Taken together, these data demonstrate that DMHCA enhances hematopoietic stem cell signaling and improves erythrocyte differentiation in diabetes. Vascular insufficiency and peripheral ischemia are known complications of diabetes. Interestingly, the first cell type that was discovered to display enhanced membrane rigidity in diabetes was erythrocytes, leading to reduced deformability, which is required for traversing small capillary lumens (55). Thus, DMHCA may correct erythrocyte dysfunction in T2D and thereby promote oxygen delivery to peripheral tissues.

Using a multi-system approach, this study demonstrates the broad and pleiotropic effects of DMHCA in the treatment of diabetes. The beneficial effects of DMHCA reported herein extend to multiple tissues and are sufficient to correct the retina-BM axis. Overall, these findings support the exciting potential of DMHCA to be used as a clinical intervention to correct a broad range of abnormalities induced by diabetes.

## Materials and Methods

### Experiment design

Male B6.BKS(D)-Lepr^db^/J (stock number:000697) Homozygous Lepr^db/db^ were diabetic and heterozygous Lepr^db/m^ were used as controls (denoted as db/db and db/m thereafter). All mice were obtained from The Jackson Laboratory (Bar Harbor, ME) and housed in the institutional animal care facilities at the University of Alabama at Birmingham (IACUC # 20917) with strict 12h:12h light: dark cycle; Animals were considered as diabetic and used in the DMHCA treatment if the serum glucose level was above 250 mg/dL on two consecutive measurements. Animals have been randomly assigned to experimental groups. The animals received DMHCA (Avanti Polar Lipidis) in their chow 8mg/kg body weight /day or base diet (5015). The animals were fed the test diets for 6 months after diabetic onset. The db/m and db/db mice were each divided in two subgroups with half the mice in each group receiving control chow and the other half DMHCA containing chow. Glycated hemoglobin was measured using the A1CNow^+^ kit (Bayer HealthCare, Sunnyvale, CA) each three months prior to euthanasia.

### Lipid extraction

Mouse retinas were subjected to monophasic lipid extraction in methanol: chloroform: water (2 : 1 : 0.74, v : v : v) as previously described (56). During lipid extraction, each sample was spiked with 100 nanograms of synthetic 19-hydroxycholesterol obtained from Steraloids (Newport, RI) for quantitation of sterols and oxysterols. Lipid extracts were resuspended using 200 µL/retina in methanol containing 0.01% butylated hydroxytoluene.

### Analysis of Free and Total Sterol Content

Sterols and oxysterols were analyzed by high resolution/accurate mass LC-MS using a Shimadzu Prominence HPLC equipped with an in-line solvent degassing unit, autosampler, column oven, and two LC-20AD pumps, coupled to a Thermo Scientific LTQ-Orbitrap Velos mass spectrometer. Lipid extracts were used directly for analysis of ‘free’ sterols and oxysterols, or subjected to alkaline hydrolysis of sterol esters for analysis of total cellular sterols as previously described (57). Gradient conditions, peak finding, and quantitation of sterols and oxysterols were performed as previously described (58). All sterol and oxysterol identifications were performed by comparison of retention time, exact mass, and MS/MS profiles to authentic standards purchased from Steraloids.

### Acellular capillaries

Eyes were fixed in 2% formalin and trypsin digest was performed for analysis of acellular capillaries as previously published (59).

### Electroretinogram (ERG)

For full-field ERG recordings, mice were dark adapted for 12 h. In preparation for the ERGs, the mice were anesthetized with intramuscular (IM) injections of ketamine and xylazine. The pupils were dilated with 1% atropine sulfate and 2.5% phenylephrine hydrochloride ophthalmic solution which also reduced sensitivity of the eyes to touch. When anesthetized, the mice were placed on a stand in a LED Ganzfeld stimulator (LKC Technologies, Gaithersburg, MD). A drop of Goniotaire solution (from Altaire pharmaceuticals and contains 2.5% hypromellose solution) was applied to each eye to ensure a good electrical connection between the electrodes and the eyes. Animals were kept on a heating pad (37 °C) during the procedure. Full-field ERGs were recorded from both eyes using the LKC system. Corneal electrodes with contact lenses were placed on each eye, and a steel subdermal needle served as the reference electrode. For grounding, a steel needle was placed in the tail. Mice were dark adapted overnight for 12 hours prior to the start of the experiment. To test scotopic responses, mice received a series of flashes at intensities −20db, −10db and 0db.

### Bone marrow analysis

Bones were harvested from mice in sterile conditions and the supernatant was washed with PBS (1x) containing a protease inhibitory cocktail (AEBSF 1mM, Aprotinin 800nM, Bestatin 50µM, E64 15µM, Leupeptin 20µM, Pepstatin A 10µM) (Thermo Scientific #78438). After centrifugation at 300 x g for 10 min at 4°, the BM supernatant was removed and the cells destined for cell culture analysis or freeze to posterior analysis. The BM supernatant was concentrated with Amicon Ultra-15 (#UFC900324) for 60 min at 3220 x g. The supernatant was used for quantification of cytokines and chemokines.

### Cytokines quantification

Levels of TNF-alpha and CCL-2 were measured on the bone marrow supernatant using Cytometric Bead Array (TNF #888299; CCL-2 #558342), and IL-3 measurement by ELISA kit (R&D system #M3000). The concentrated supernatant was incubated with beads and acquired on a BD FACSCelesta following the manufacturer’s instructions. The concentration was determined using a standard curve and analyzing on BD FACSArray software. Protein assay was performed to normalize the concentration in pg/mg.

### Colony-forming unit

Analysis of colony-forming units was performed as previously published (60). BM cells (10^5^) were aliquot in 1 mL of Iscove’s modified Dulbecco’s medium (StemCell Technologies #07700) then 300 µL of this solution was added in 3 mL of MethoCult ^TM^ GF3434 (STEMCELL Technologies) to obtain a final concentration of 10^4^ cells. The cell suspension (1.1 mL) was seed in 35-mm dishes. Culture were placed in the incubator 37 °C with 5% of CO_2_ for 12 days, following of counting of colonies based on the morphology of cells based on manufacturer’s instructions. Counting was performed in duplicate.

### qRT-PCR

RNA was isolated according with the RNeasy mini kit (74106; Qiagen, Valencia, CA) according to manufacturer’s instructions. First-strand complementary DNA was synthesized from isolated RNA using iScript II reverse transcription supermix (1708841; Bio-Rad). Prepared cDNA was mixed with SsoAdvanced Universal SYBR Green Mix (172570; Bio-Rad) and sets of gene-specific forward and reverse primers (ABCA1, CCL-2) (Bio-Rad) and subjected to real-time PCR quantification using the CFX384 Real Time PCR. All reactions were performed in triplicate. Cyclophilin A was used as a control, and results were analyzed using the comparative Ct method and Ct values were normalized to Cyclophilin A levels. Data is shown as normalized relative to control levels or as non-normalized raw expression levels.

### Flow cytometry analysis

2 x 10^6^ cells isolated from bone marrow and 100 µL of peripheral blood were incubated with Ammonium Chloride solution (Stemcell Tecnologies #07850) for 14 min on ice to lyse red cells. The cells were washed twice with PBS 2% FBS and incubated with a cocktail of primary antibodies (viability dye 510, Ly6G, CD45, CD11b, CCR2, CD31, CD133, Ly6C and Flk-1) for 30 minutes at 4°C in the dark (for the panel of antibodies for myeloid analysis refer to supplemental table 2). To quantify precursor cells in the bone marrow, 1 x 10^6^ cells were incubate with a cocktail of antibodies containing c-Kit (CD117), viability dye 510, FcyRII/III, Sca-1, Lineage cocktail, CD34, FLT3 (CD135) and CD127 (for the panel of antibodies for precursor analysis refer to supplemental table 3). After washing, the cells were acquired on BD FACSCelesta. Retina cells were isolated by incubating the entire retina with the digesting buffer (RPMI 5% FBS, 10 µg/mL collagenase D and 300µ/ml Dnase) for 1 hour at 37 °C. The suspension of cells were filtrated in a 40 µm cell strainer and incubated with a cocktail of antibodies containing anti-F4/80, Viability dye 510, Ly6G, CD45, CD11b, CCR2, CD133, CD206, Ly6C and Flk-1 (for the list of antibodies, refer to supplemental table 4). All the flow cytometry analyses were performed using FlowJo software.

### 10X Genomics single-cells

For single-cell RNA-Seq (scRNA-seq), BM single-cell suspensions were generated from 9 month old untreated and DMHCA treated db/db mice. Briefly, BM cells were aspirated from femur samples and filtered through #40 µm mesh. Single-cell suspensions were column enriched for Sca-1^+^, FACS sorted using lineage^-^ and c-Kit1^+^ markers, and assayed for viability using trypan blue. Viable cells were then run through the 10X genomics platform for droplet-based single-cell barcoding and cDNA generation. Illumina HiSeq was used for cDNA sequencing. The 10X Genomics software Cell Ranger (version 3.1.0) was used for quality control of sequencing reads, FASTQ file generation, and demultiplexing. The STAR software was used for read alignment using the mouse mm10 reference genome.

### Single-cell RNA-seq data analysis

For scRNA-seq data analysis, multiple software platforms were used including Scanpy, Monocle, and Seurat. The authors are grateful for Luecken and Theis (61) whose tutorial on the current best practices in single-cell RNA-seq analyses formed the foundation of the analysis herein. For transcript quality control, Scanpy was used to plot histograms of total counts per cell and genes per cell, which were then used to identify cutoffs that eliminated doublets and damaged cells. Additionally, a mitochondrial gene percentage cutoff of 20% was used to further eliminate damaged cells. After quality control, 5,103 cells were recovered from the untreated diabetic group and 5,152 cells from the DMHCA treated group, for a total of 10,255 cells. The two treatment groups were then concatenated to form a single adata file, and a minimum cutoff of 10 cells per gene was used to eliminate lowly represented genes. Normalization was performed using Scran, which employs a pooling-based size factor estimation method to normalize single-cell transcript data across heterogenous cell populations (62). Scanpy was then used to perform complete cell cycle regression using the cell cycle genes identified by Tirosh et al (63). ComBat was used for batch correction with cell cycle scores included as covariates (64). Scanpy was then used to select 4,500 highly variable genes, and the UMAP plot was generated using a resolution of 0.8. Subclustering was then performed to arrive at the final UMAP representation of 13 clusters. Scanpy was used to identify marker genes for each cluster. The assignment of cluster identities was guided by the expression of lineage-specific marker genes identified in previous scRNA-seq datasets on murine HSCs (44, 45, 65). In the case of novel undefined cell populations, clusters were identified based on their unique expression patterns and the presence/absence of lineage-specific markers. Compositional analysis was done using Scanpy.

To identify differentially expressed genes across total samples or specific cluster subpopulations, MAST was used and the analysis included the technical covariate number of genes expressed per cell (66). An FDR cutoff of 0.01 was used in the total sample DGE while a cutoff of 0.05 was used in the subpopulation analyses. Pathway enrichment analysis was performed using Ingenuity Pathways Analysis (IPA), with all canonical pathways used in the total sample DGE and secondary messenger/intracellular signaling pathways in the subpopulation analyses.

Pseudotime trajectory analysis was performed using Monocle with the naïve B cell population removed and max components set to 3. PAGA analysis was performed using the preprocessed adata file with the threshold set to 0.07. Seurat was used to graph the violin plots.

### Immunofluoresence staining

Enucleated eyes were fixed in 4% paraformaldehyde overnight at 4 °C. The next day, eyes were washed 2X for 5 min with PBS(1x) follow by incubation in 30% sucrose for 48 hrs and then snap frozen in optical cutting temperature (OCT) compound. Retinal crossections (12 µm) were processed for immunostaining using the following antibodies: rat monoclonal to CD45 antibody (clone 30-F11, R&D Systems, 1:50). Sections were pre-incubated with 5% goat serum (Invitrogen) in PBS for 1hr, followed by incubation with primary antibodies (in 1% normal goat serum) overnight at 4°C. Alexa Fluor 594 was used as the secondary antibody. Positive cells were counted from 3-5 sections at 100 µm interval for each eye with a minimum of four images per section. Retinal sections were imaged using a confocal scanning laser microscope (ZEISS LSM 700 confocal microscope system with Axio Observer; Carl Zeiss Mditec, Jena, Germany) and the images were analyzed by using Zen lite software for colocalization analysis.

### Membrane fluidity

CAC cells isolated from control or diabetic human subjects or mouse models were left untreated or treated with DMHCA at 10 µM overnight (16-18hrs) in StemSpan SFEM + CC 110 media at 37 °C.

Perylene stock solution (1 mM) was prepared in dimethyl sulfoxide DMSO (Sigma Aldrich), aliquoted and stored in a −80°C freezer. CACs were stained with 10µM perylene for 10 min, centrifuged (0.8 or 1 x 10^3^ G, 10 min, 20 °C, Fisher Scientific AccuSpin Micro 17R) to remove access dye and kept at 37 °C until use. A 0.8-μL aliquot was pipetted on a microscope slide (Thermo Scientific plain microscope slides, precleaned, 25 x 75 mm x 1 mm thick) and a cover-glass was placed on top (Corning, 22 x 22 mm of 1 mm thickness). The assembly was turned upside down and positioned on the flat stage of the FRAP instrument (*vide infra*).

### Fluorescence Recovery after Photobleaching (FRAP) Measurements

FRAP was performed as previously described (67, 68).

Briefly, samples were placed on a motorized stage of a confocal scanning microscope (Nikon C2+), and images using 40X objective and 405 excitation laser. NIS-Elements Acquisition imaging software (v 4.30) was used for FRAP experiments, with settings of pixel dwell 1.9, size 512, normal, DAPI checked, HV between 90-145 (typically ∼125), offset of 10, and laser power 0.71. The 40X objective was used to locate cells and region of interest (ROI) was either 3 or 2 μm diameter for human or mouse samples, respectively, one used as the stimulating spot and the other as a standard, placed in a dark, non-fluorescent spot. A continuous scanning time measurement was performed to ensure the cell did not move, indicated by a constant fluorescent intensity value over time. There was 1 minute of data acquisition (for 61 loops) at 1 sec intervals, bleaching for 1 second (4 loops) with no delay in intervals, and acquisition for another 5 min (301 loops) at 1 sec intervals. Data were fit using IGOR Pro software (WaveMetrics Inc.).

### Statistical analysis

All values are expressed as mean ± SEM. A value of p < 0.05 was considered to be statistically significant. Statistical tests were performed using statistics software (GraphPad Software; La Jolla, CA). Mann-Whitney test t was used for comparisons between two groups.

## Acknowledgements

This study was supported by the National Institutes of Health Grants R01EY025383, R01EY012601, R01EY028858, R01EY028037, to M.B. Grant; R01EY030766, R01EY016077 to J.V. Busik F32EY028426 to S. Hammer; T32HL134640-01 to M. Dupont; T32HL105349 to J.L. Floyd.

## References

1. The ACCORD Study Group and ACCORD Eye Study Group. Effects of Medical Therapies on Retinopathy Progression in Type 2 Diabetes. N Engl J Med. 2010;363(3):233–244. doi:10.1056/NEJMoa1001288

2. Sandhoff R, Brügger B, Jeckel D, et al. Determination of cholesterol at the low picomole level by nano-electrospray ionization tandem mass spectrometry. J Lipid Res. 1999;40(1):126–132.

3. Stancu C, Sima A. Statins: mechanism of action and effects. J Cellular Mol Med. 2001;5(4):378–387. doi:10.1111/j.1582-4934.2001.tb00172.x

4. Niesor EJ, Schwartz GG, Perez A, et al. Statin-Induced Decrease in ATP-Binding Cassette Transporter A1 Expression via microRNA33 Induction may Counteract Cholesterol Efflux to High-Density Lipoprotein. Cardiovasc Drugs Ther. 2015;29(1):7–14. doi:10.1007/s10557-015-6570-0

5. Zheng W, Mast N, Saadane A, et al. Pathways of cholesterol homeostasis in mouse retina responsive to dietary and pharmacologic treatments. J Lipid Res. 2015;56(1):81–97. doi:10.1194/jlr.M053439

6. Bryszewska M, Watala C, Torzecka W. Changes in fluidity and composition of erythrocyte membranes and in composition of plasma lipids in Type I diabetes. Br J Haematol. 1986;62(1):111–116. doi:10.1111/j.1365-2141.1986.tb02906.x

7. Kruit JK, Wijesekara N, Fox JEM, et al. Islet Cholesterol Accumulation Due to Loss of ABCA1 Leads to Impaired Exocytosis of Insulin Granules. Diabetes. 2011;60(12):3186–3196. doi:10.2337/db11-0081

8. Simons K, Ehehalt R. Cholesterol, lipid rafts, and disease. J Clin Invest. 2002;110(5):597-603. doi:10.1172/JCI0216390

9. Chen W, Jump DB, Esselman WJ, et al. Inhibition of cytokine signaling in human retinal endothelial cells through modification of caveolae/lipid rafts by docosahexaenoic acid. Invest Ophthalmol Vis Sci. 2007 Jan;48(1):18–26

10. Wang B, Tontonoz P. Liver X receptors in lipid signalling and membrane homeostasis. Nat Rev Endocrinol. 2018;14(8):452–463. doi:10.1038/s41574-018-0037-x

11. Janowski BA, Willy PJ, Devi TR, et al. An oxysterol signalling pathway mediated by the. Nature. 1996;383(6602):728–731. doi:10.1038/383728a0

12. Venkateswaran A, Laffitte BA, Joseph SB, et al. Control of cellular cholesterol efflux by the nuclear oxysterol receptor LXRalpha. Proceedings of the National Academy of Sciences. 2000;97(22):12097–12102. doi:10.1073/pnas.200367697

13. Rong X, Albert CJ, Hong C, et al. LXRs Regulate ER Stress and Inflammation through Dynamic Modulation of Membrane Phospholipid Composition. Cell Metabolism. 2013;18(5):685–697. doi:10.1016/j.cmet.2013.10.002

14. Cermenati G, Abbiati F, Cermenati S, et al. Diabetes-induced myelin abnormalities are associated with an altered lipid pattern: protective effects of LXR activation. J Lipid Res. 2012;53(2):300–310. doi:10.1194/jlr.M021188

15. Cermenati G, Giatti S, Cavaletti G, et al. Activation of the Liver X Receptor Increases Neuroactive Steroid Levels and Protects from Diabetes-Induced Peripheral Neuropathy. J Neurosci. 2010;30(36):11896–11901. doi:10.1523/JNEUROSCI.1898-10.2010

16. Joseph SB, McKilligin E, Pei L, et al. Synthetic LXR ligand inhibits the development of atherosclerosis in mice. Proceedings of the National Academy of Sciences. 2002;99(11):7604–7609. doi:10.1073/pnas.112059299

17. Hazra S, Rasheed A, Bhatwadekar A, et al. Liver X Receptor Modulates Diabetic Retinopathy Outcome in a Mouse Model of Streptozotocin-Induced Diabetes. Diabetes. 2012;61(12):3270–3279. doi:10.2337/db11-1596

18. Hammer SS, Beli E, Kady N, et al. The Mechanism of Diabetic Retinopathy Pathogenesis Unifying Key Lipid Regulators, Sirtuin 1 and Liver X Receptor. EBioMedicine. 2017;22:181-190. doi:10.1016/j.ebiom.2017.07.008

19. Chisholm JW, Hong J, Mills SA, Lawn RM. The LXR ligand T0901317 induces severe lipogenesis in the db/db diabetic mouse. J Lipid Res. 2003;44(11):2039–2048. doi:10.1194/jlr.M300135-JLR200

20. Kirchgessner TG, Sleph P, Ostrowski J, et al. Beneficial and Adverse Effects of an LXR Agonist on Human Lipid and Lipoprotein Metabolism and Circulating Neutrophils. Cell Metabolism. 2016;24(2):223–233. doi:10.1016/j.cmet.2016.07.016

21. Quinet EM, Savio DA, Halpern AR, Chen L, Miller CP, Nambi P. Gene-selective modulation by a synthetic oxysterol ligand of the liver X receptor. J Lipid Res. 2004;45(10):1929–1942. doi:10.1194/jlr.M400257-JLR200

22. Kratzer A1, Buchebner M, Pfeifer T, et al., Synthetic LXR agonist attenuates plaque formation in apoE-/- mice without inducing liver steatosis and hypertriglyceridemia. J Lipid Res. 2009;50(2):312–26. doi: 10.1194/jlr.M800376-JLR200

23. Fessler MB. The challenges and promise of targeting the Liver X Receptors for treatment of inflammatory disease. Pharmacol Ther. 2018;181:1–12. doi: 10.1016/j.pharmthera.2017.07.010

24. Pfeifer T, Buchebner M, Chandak PG, et al. Synthetic LXR agonist suppresses endogenous cholesterol biosynthesis and efficiently lowers plasma cholesterol. Curr Pharm Biotechnol. 2011;12(2):285–292. doi:10.2174/138920111794295774

25. Repa JJ. Regulation of mouse sterol regulatory element-binding protein-1c gene (SREBP-1c) by oxysterol receptors, LXRalpha and LXRbeta. Genes & Development. 2000;14(22):2819–2830. doi:10.1101/gad.844900

26. Schulman IG. Liver X receptors link lipid metabolism and inflammation. FEBS Lett. 2017;591(19):2978–2991. doi:10.1002/1873-3468.12702

27. Song X, Wu W, Gabbi C, et al. Retinal and optic nerve degeneration in liver X receptor β knockout mice. Proc Natl Acad Sci USA. 2019;116(33):16507–16512. doi:10.1073/pnas.1904719116

28. Choudhary M, Ismail EN, Yao PL, et al., LXRs regulate features of age-related macular degeneration and may be a potential therapeutic target. JCI Insight. 2020 16;5(1). pii: 131928. doi: 10.1172/jci.insight.131928.

29. Ito A, Hong C, Rong X, et al. LXRs link metabolism to inflammation through Abca1-dependent regulation of membrane composition and TLR signaling. eLife. 2015;4:e08009. doi:10.7554/eLife.08009

30. Duan Y, Prasad R, Feng D, et al. Bone Marrow-Derived Cells Restore Functional Integrity of the Gut Epithelial and Vascular Barriers in a Model of Diabetes and ACE2 Deficiency. Circ Res. 2019;125(11):969–988. doi:10.1161/CIRCRESAHA.119.315743

31. Dimmeler S, Zeiher AM. Vascular repair by circulating endothelial progenitor cells: the missing link in atherosclerosis? J Mol Med. 2004;82(10):671–677. doi:10.1007/s00109-004-0580-x

32. Fadini GP, de Kreutzenberg S, Agostini C, et al. Low CD34+ cell count and metabolic syndrome synergistically increase the risk of adverse outcomes. Atherosclerosis. 2009;207(1):213–219. doi:10.1016/j.atherosclerosis.2009.03.040

33. Jarajapu YPR, Hazra S, Segal M, et al. Vasoreparative Dysfunction of CD34+ Cells in Diabetic Individuals Involves Hypoxic Desensitization and Impaired Autocrine/Paracrine Mechanisms. Madeddu P, ed. PLoS ONE. 2014;9(4):e93965. doi:10.1371/journal.pone.0093965

34. Li Calzi S, Shaw LC, Moldovan L, et al. Progenitor cell combination normalizes retinal vascular development in the oxygen-induced retinopathy (OIR) model. JCI Insight. 2019;4(21):e129224. doi:10.1172/jci.insight.129224

35. Rasheed A, Tsai R, Cummins CL. Loss of the Liver X Receptors Disrupts the Balance of Hematopoietic Populations, With Detrimental Effects on Endothelial Progenitor Cells. JAHA. 2018;7(10). doi:10.1161/JAHA.117.007787

36. Laffitte BA1, Joseph SB, Walczak R, et al. Autoregulation of the human liver X receptor alpha promoter. Mol Cell Biol. 2001;21(22):7558–68

37. Blanchard GJ, Busik JV. Interplay between Endothelial Cell Cytoskeletal Rigidity and Plasma Membrane Fluidity. Biophys J. 2017;14;112(5):831-833. doi: 10.1016/j.bpj.2017.01.013.

38. Chakravarthy H, Navitskaya S, O’Reilly S, et al. Role of Acid Sphingomyelinase in Shifting the Balance Between Proinflammatory and Reparative Bone Marrow Cells in Diabetic Retinopathy. Stem Cells. 2016;34(4):972–83. doi: 10.1002/stem.2259.

39. Ayee MAA, LeMaster E, Shentu TP, et al. Molecular-Scale Biophysical Modulation of an Endothelial Membrane by Oxidized Phospholipids. Biophys J. 2017;24;112(2):325-338. doi: 10.1016/j.bpj.2016.12.002.

40. Bogdanov P, Corraliza L, Villena JA, et al. The db/db mouse: a useful model for the study of diabetic retinal neurodegeneration. PLoS One. 2014;16;9(5):e97302. doi: 10.1371/journal.pone.0097302

41. Cheung AK, Fung MK, Lo AC, et al. Aldose reductase deficiency prevents diabetes-induced blood-retinal barrier breakdown, apoptosis, and glial reactivation in the retina of db/db mice. Diabetes. 2005;54(11):3119–25

42. Ferraro F, Lymperi S, Méndez-Ferrer S, et al. Diabetes impairs hematopoietic stem cell mobilization by altering niche function. Sci Transl Med. 2011;12;3(104):104ra101. doi: 10.1126/scitranslmed.3002191

43. Hazra S, Jarajapu YP, Stepps V, et al. Long-term type 1 diabetes influences haematopoietic stem cells by reducing vascular repair potential and increasing inflammatory monocyte generation in a murine model. Diabetologia. 2013;56(3):644–53. doi:10.1007/s00125-012-2781-0

44. Tusi BK, Wolock SL, Weinreb C, et al. Population snapshots predict early haematopoietic and erythroid hierarchies. Nature. 2018;555(7694):54–60. doi: 10.1038/nature25741

45. Dahlin JS, Hamey FK, Pijuan-Sala B. A single-cell hematopoietic landscape resolves 8 lineage trajectories and defects in Kit mutant mice. Blood. 2018;131(21):e1–e11. doi: 10.1182/blood-2017-12-821413

46. Pehkonen P, Welter-Stahl L, Diwo J, et al. Genome-wide landscape of liver X receptor chromatin binding and gene regulation in human macrophages. BMC Genomics. 2012; 31;13:50. doi: 10.1186/1471-2164-13-50

47. Bahrami S, Drabløs F. Gene regulation in the immediate-early response process. Adv Biol Regul. 2016;62:37–49. doi: 10.1016/j.jbior.2016.05.001

48. Okada S, Fukuda T, Inada K, et al. Prolonged expression of c-fos suppresses cell cycle entry of dormant hematopoietic stem cells.Blood. 1999 Feb 1;93(3):816–25

49. Okada S, Wang ZQ, Grigoriadis AE, et al. Mice lacking c-fos have normal hematopoietic stem cells but exhibit altered B-cell differentiation due to an impaired bone marrow environment. Mol Cell Biol. 1994;14(1):382–90

50. Spencer JA, Ferraro F, Roussakis E, et al. Direct measurement of local oxygen concentration in the bone marrow of live animals. Nature. 2014;10;508(7495):269–73. doi: 10.1038/nature13034

51. Rabbani PS, Soares MA, Hameedi SG, et al. Dysregulation of Nrf2/Keap1 Redox Pathway in Diabetes Affects Multipotency of Stromal Cells. Diabetes. 2019;68(1):141–155. doi: 10.2337/db18-0232

52. Basu P, Morris PE, Haar JL, et al. KLF2 is essential for primitive erythropoiesis and regulates the human and murine embryonic beta-like globin genes in vivo. Blood. 2005;106: 2566–2571

53. Matsumoto N, Kubo A, Liu H, et al. Developmental regulation of yolk sac hematopoiesis by Kruppel-like factor 6. Blood. 2006;107(4):1357–65. Epub 2005

54. Mayers S, Moço PD, Maqbool T, et al. Establishment of an erythroid progenitor cell line capable of enucleation achieved with an inducible c-Myc vector. BMC Biotechnol. 2019;19(1):21. doi: 10.1186/s12896-019-0515-9

55. McMillan DE, Utterback NG, La Puma J. Reduced erythrocyte deformability in diabetes.Diabetes. 1978;27(9):895–901

56. Lydic TA, Busik JV, Reid GE. A monophasic extraction strategy for the simultaneous lipidome analysis of polar and nonpolar retina lipids. J Lipid Res. 2014;55(8):1797–809. doi: 10.1194/jlr.D050302.

57. McDonald JG, Thompson BM, McCrum EC et al. Extraction and analysis of sterols in biological matrices by high performance liquid chromatography electrospray ionization mass spectrometry. Methods Enzymol. 2007;432:145–70

58. Machacek M, Saunders H, Zhang Z, et al.Elevated O-GlcNAcylation enhances pro-inflammatory Th17 function by altering the intracellular lipid microenvironment. J Biol Chem. 2019;294(22):8973–8990. doi: 10.1074/jbc.RA119.008373

59. Bhatwadekar A, Glenn JV, Figarola JL, et al. A new advanced glycation inhibitor, LR-90, prevents experimental diabetic retinopathy in rats. Br J Ophthalmol 2008;92:545–547

60. Duan Y, Beli E, Li Calzi S, et al. Loss of Angiotensin-Converting Enzyme 2 Exacerbates Diabetic Retinopathy by Promoting Bone Marrow Dysfunction. Stem Cells. 2018;36(9):1430–1440. doi: 10.1002/stem.2848

61. Luecken MD, Theis FJ. Current best practices in single-cell RNA-seq analysis: a tutorial. Mol Syst Biol. 2019;15(6):e8746. doi: 10.15252/msb.20188746.

62. Lun AT, Bach K, Marioni JC. Pooling across cells to normalize single-cell RNA sequencing data with many zero counts. Genome Biol. 2016 27;17:75. doi: 10.1186/s13059-016-0947-7

63. Tirosh I, Izar B, Prakadan SM, et al. Dissecting the multicellular ecosystem of metastatic melanoma by single-cell RNA-seq. Science. 2016;352(6282):189–96. doi:10.1126/science.aad0501

64. Johnson WE, Li C, Rabinovic A. Adjusting batch effects in microarray expression data using empirical Bayes methods. Biostatistics. 2007;8(1):118–27

65. Han X, Wang R, Zhou Y. Mapping the Mouse Cell Atlas by Microwell-Seq. Cell. 2018;172(5):1091–1107.e17. doi: 10.1016/j.cell.2018.02.001

66. Finak G, McDavid A, Yajima M et al. MAST: a flexible statistical framework for assessing transcriptional changes and characterizing heterogeneity in single-cell RNA sequencing data. Genome Biol. 2015;10(16):278. doi: 10.1186/s13059-015-0844-5

67. Axelrod D, Koppel DE, Schlessinger J, Elson E, Webb WW. Mobility measurement by analysis of fluorescence photobleaching recovery kinetics. Biophys J. 1976;16(9):1055–69

68. Ishikawa-Ankerhold H, Ankerhold R, Drummen G. Fluorescence Recovery After Photobleaching (FRAP). Encyclopedia of Life Sciences. John Wiley & Sons, Ltd: Chichester 2014, 1-11.

