## Supplemental file for "Selective LXR agonist, DMHCA, corrects the retina-bone marrow axis in type 2 diabetes"

| **Subject ID** | **Age** | **Type** | **HGBA1C** | **Blood Sugar** | **Drugs** | **Left Eye** | **Right Eye** |
| --- | --- | --- | --- | --- | --- | --- | --- |
| **Control** | | | | | | | |
| DYS 019 | 49 | Control | Control | Control | Control | Control | Control |
| DYS 020 | 50 | Control | Control | Control | Control | Control | Control |
| DYS 025 | 48 | Control | Control | 91 | Control | Control | Control |
| DYS 027 | 24 | Control | Control | 77 | Control | Control | Control |
| DYS 028 | 68 | Control | Control | Control | Control | Control | Control |
| DYS 029 | 21 | Control | Control | Control | Control | Control | Control |
| DYS 045 | 69 | Control | Control | 34 | Control | Control | Control |
| LXR-007 | 28 | Control | Control | Control | Control | Control | Control |
| LXR-008 | 29 | Control | Control | Control | Control | Control | Control |
| LXR-016 | 42 | Control | Control | Control | Control | Control | Control |
| LXR-017 | 26 | Control | Control | Control | Control | Control | Control |
| LXR-019 | 53 | Control | Control | Control | Control | Control | Control |
| LXR-026 | 50 | Control | 4.8 | 94 | Control | Control | Control |
| LXR-030 | 48 | Control | Control | 91 | Control | Control | Control |
| LXR-033 | 64 | Control | Control | 64 | Control | Control | Control |
| LXR-034 | 50 | Control | Control | Control | Control | Control | Control |
| LXR-035 | 55 | Control | Control | Control | Control | Control | Control |
| LXR-036 | 29 | Control | Control | 59 | Control | Control | Control |
| LXR 048 | 71 | Control | 4.9 | 33 | Control | Control | Control |
| **Type 2** | | | | | | | |
| DYS 013 | 52 | Type 2 | 5.5 | 140 | Aspir 81, Carisoprodol, Clotrimazole-Betamethasone, Fenofibrate Nanocrystallized, Immune Support Complex, Lipitor, Metformin, Niacin ER, Novolog Pen Fill 100, Rosuvastatin, Valsartan, Vitamin D3, Indapamide | None | None |
| DYS 016 | 40 | Type 2 | Unknown | Unknown | Apidra, Avasin, Lantus | Proliferative | Proliferative |
| DYS 017 | 63 | Type 2 | 8.1 | 229 | Fibercon, Losartan, Novofine , Lortab, Trazadone, Oxycodone, Meloxicam, Humalog Mix, Hydrochlorothiazide, Multivitamins, Metformin, Tramadol, Aspirin, Primidone, Simvastatin , Percocet, Bupropion | None | None |
| DYS 018 | 58 | Type 2 | Unknown | Unknown | Amlodipine Besylate ,Lasix, Losartan Potassium, Metoprolol Tartrate, Paxil, Ranitidine HCI | Proliferative | Proliferative |
| DYS 022 | 65 | Type 2 | 7.1 | 136 | Actos, Alprazolam, Fluticasone, Nasal Spray, Suspension, Furosemide, Furosemide,  Gabapentin, Humalog Mix Insulin, Hydrocodone, Acetaminophen,  Jardiance, Kionex,  Levothyroxine, Lisinopril,  Pramipexole, Primidone, Silver Sulfadiazine, Simvastatin,  Spironolactone, Sulfamethoxazole,  Tizanidine, Tramadol, Venlafaxine, Victoza,  Vitamin D2 | Mild Nonprliferative | Mild Nonprliferative |
| DYS 023 | 77 | Type 2 | Unknown | 88 | Finasteride, Hydrocodone, Metformin, Methimazole, Lisinopril, Nitrofurantoin Monohydrate, Oxybutynin Chloride ER, Sulfamethoxazole, Trazodone, Oxycodone- acetaminophen, Warfarin | None | None |
| DYS 024 | 50 | Type 2 | Unknown | 331 | Unknown | None | None |
| LXR-009 | 52 | Type 2 | 5.5 | 150 | Aspir-81( Delayed Release), Carisoprodol, Clotrimazole-Betamethasone Cream ,Fenofibrate Nanocrystallized, Immune Support Complex, Indapamide, Lipitor, Metformin, Niacin, Novolog Penfill, Rosuvastatin, Valsartan, Vitamin D3,Zoloft | None | None |
| LXR-010 | 66 | Type 2 | Unknown | Unknown | Unknown | None | None |
| LXR-015 | 77 | Type 2 | Unknown | 88 | Finasteride, Hydrocodone, Metformin, Lisinopril, Methimazole, Nitrofurantoin, Monohydrate, Oxybutynin, Sulfamethoxazole, Trazodone, Oxycodone, Warfarin | None | None |
| LXR-020 | 60 | Type 2 | Unknown | Unknown | Prednisolone Acetate, Tobramycin, Albuterol Sulfate, Lisinopril, Ibuprofen | Proliferative | Proliferative |
| LXR-027 | 41 | Type 2 | 6.7 | 76 | Maxitrol Drop, Acetazolamide, Amlodipine, Apidra, Atorvastatin, Avastin, Gabapentin, Hydrocodone, Lantus, Lisinopril, Novolin, Prednisone, Sumatriptan Tramadol, Trazodone | Proliferative | Proliferative |
| LXR-028 | 62 | Type 2 | 8.1 | 229 | Fibercon, Losartan, Novofine , Lortab, Trazadone, Oxycodone, Meloxicam, Humalog Mix, Hydrochlorothiazide, Multivitamins, Metformin, Tramadol, Aspirin, Primidone, Simvastatin , Percocet, Bupropion | None | None |
| LXR-029 | 54 | Type 2 | 8.4 | Unknown | Unknown | None | None |
| LXR-032 | 48 | Type 2 | Unknown | 186 | Cozaar, Lantus, Norvasc, Novolog, Simvastatin | Proliferative | Proliferative |
| LXR-043 | 52 | Type 2 | Unknown | 427 | Aspirin, Meclizine, Naprosyn, Neurontin, Novolin R, Sumatriptan | Severe NPDR | Severe NPDR |
| LXR-044 | 55 | Type 2 | 11.2 | 293 | Ergocalciferol, Metformin, Novolog Mix, Proair HFA, Simvastatin | Moderate NPDR | Moderate NPDR |
| LXR-046 | 59 | Type 2 | 6.0 | 194 | Amlodipine, Atorvastatin, Bupropion, Ciprofloxacin, Clindamycin, Gabapentin, Lisinopril 40mg, Lisinopril 20mg, Metformin, Sulfamethoxazole, Sure Comfort Insulin, Synthroid | Unspecifiied | Unspecified |
| LXR-047 | 69 | Type 2 | 7.4 | 79 | Azelastine, Amlodipine, Azithromycin, Clonidine, Enalapril, Furosemide, Metoprolol, Novolin, Potassium Chloride, Sure Comfort Insulin Syringe | Moderate NPDR | Moderate NPDR |

**Supplemental Table 1**. Identification of human subjects.

All events

Single Cells

Live Cells

CD45+ Cells

CD11b-FLK-1+

CAC

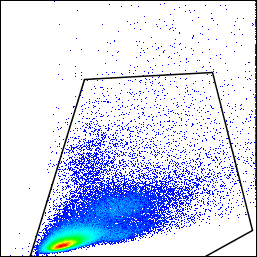

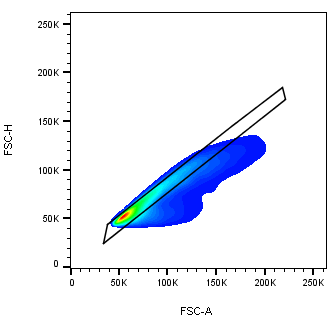

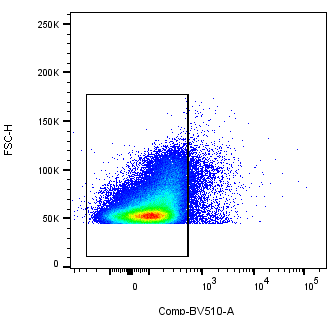

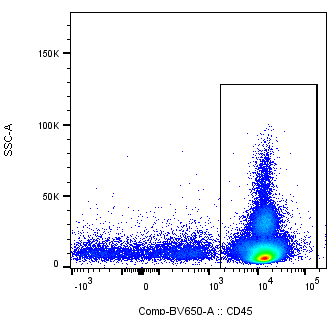

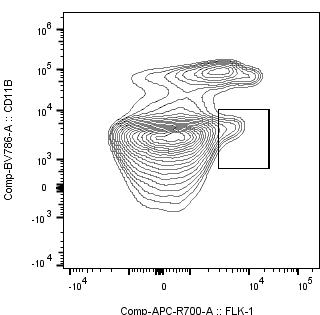

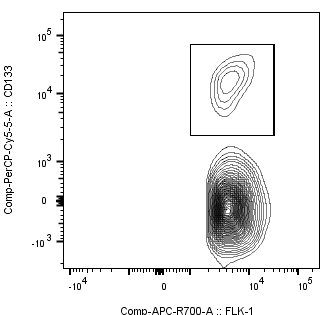

**Supplemental Figure 1 – Gate Strategy of CAC on BM cells and PBMCs.** CD45^+^ cells were gated after exclusion of debris, doublets and dead cells. The FLK-1^+^CD11b^-^ cells were defined for further selection of CD133^+^ cells.

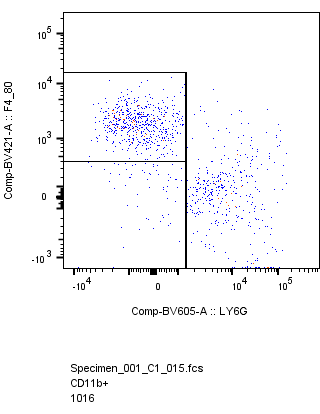

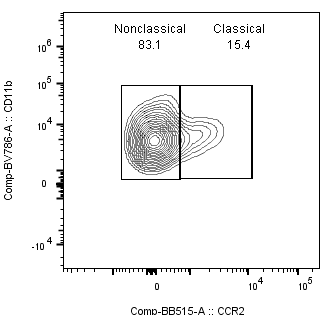

Classical

Nonclassical

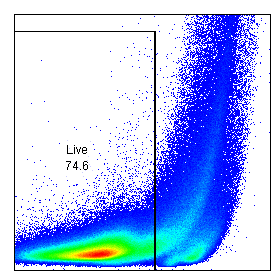

Single Cells

Live Cells

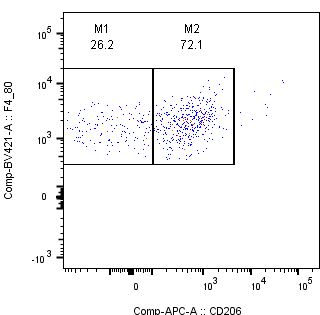

M1

M2

F4/80^+^Ly6G^-^

F4/80^-^Ly6G^-^

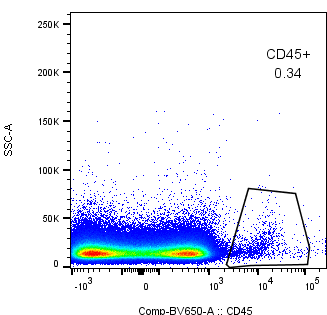

CD45^+^

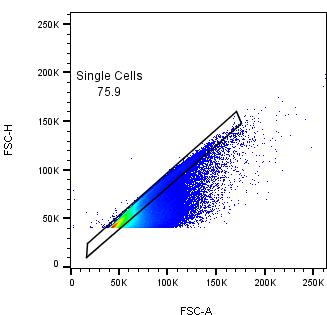

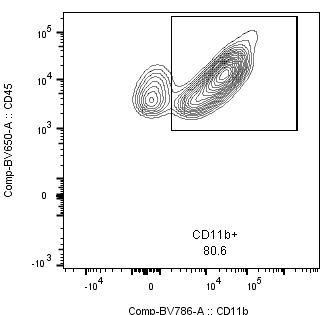

CD11b^+^

**Supplemental Figure 2 – Gate Strategy to define monocytes and macrophages subtypes on Retina**. CD45^+^ cells were gated after exclusion of doublets and dead cells. The CD11b^+^ expressing F4/80 and lacking expression of Ly6G were defined as macrophages, further the CD206^+^ cells were gated to address M2 and CD206^-^ to define M1 macrophages. Monocytes (CD11b^+^Ly6G^-^F4/80^-^) were divided into CCR2^+^ fraction comprised of classical monocytes and CCR2^-^ fr action defined as nonclassical monocytes.

All events

Single Cells

Live Cells

Monocytes

Ly6C^+^Ly6G^-^

CD11b^+^

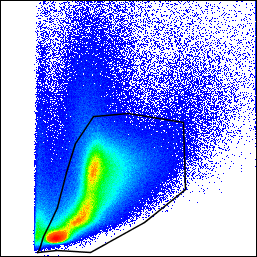

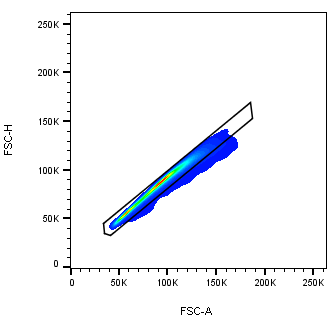

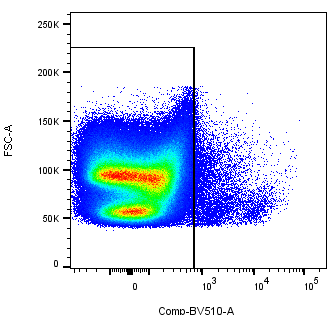

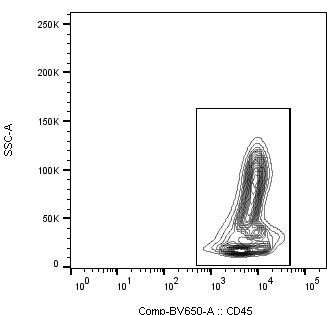

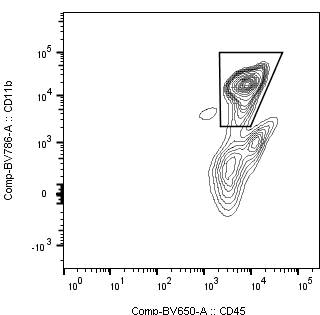

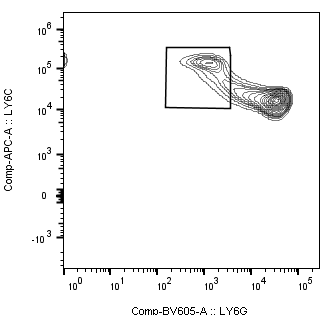

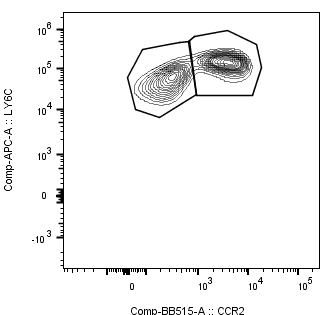

CD45^+^

**Supplemental Figure 3 – Gate Strategy of subtypes of monocytes on BM cells and PBMCs.** CD45^+^ cells were gated after exclusion of debris, doublets and dead cells. The CD11b^+^ expressing Ly6C and lacking expression of Ly6G were defined as monocytes. Monocytes were divided into CCR2^+^ fraction comprised of classical monocytes and CCR2^-^ fraction defined as nonclassical monocytes.

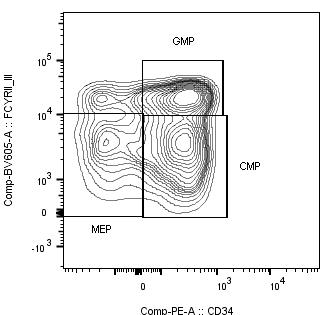

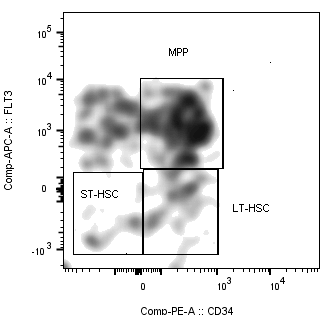

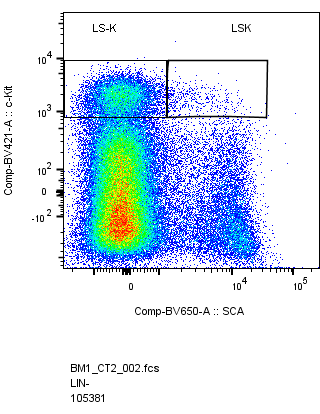

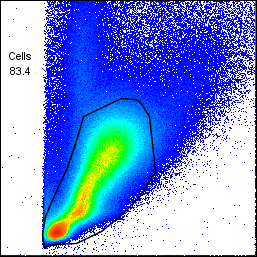

All events

BM Cells

Live Cells

Single Cells

Lineage neg

LSK

LS-K

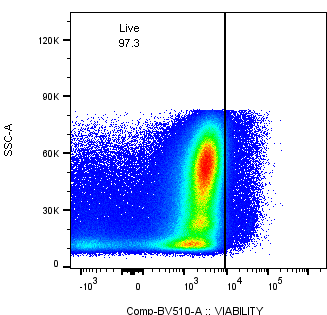

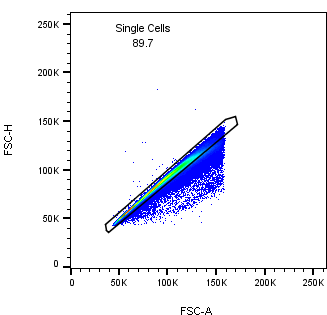

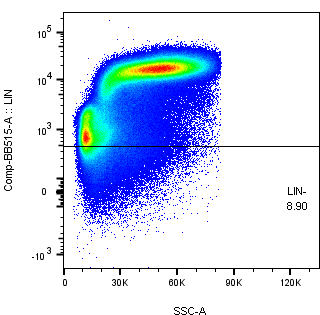

**Supplemental Figure 4 – Gate Strategy of HSC on BM.** Lin^-^ cells were gated after exclusion of debris, doublets and dead cells. Sca-1^-^c-Kit^+^ cells (LS-K) were divided in CD34^+^FcgRII/III^+^ (GMP), CD34^+^FcgRII/III^-^ (CMP) and CD34^-^FcgRII/III^-^ (MEP). Sca^+^c-Kit^+^ (LSK) were divided in CD34^+^FLT3^+^ (MPP), CD34^+^FLT3^-^ (LT-HSC) and CD34^-^FLT3^-^ (ST-HSC).

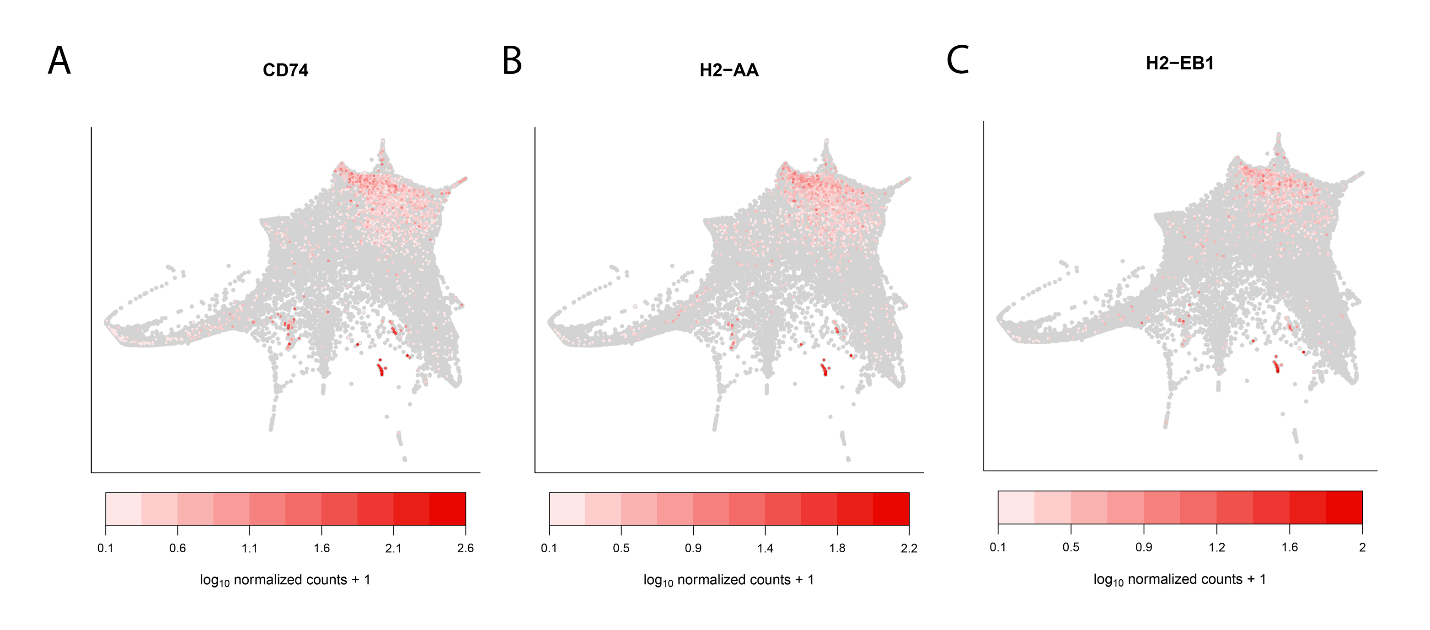

**Supplemental Figure 5. Expression of dendritic cell markers in scRNA-seq dataset from Dahlin et al 2018.** scRNA-seq was performed on 44,802 HSCs derived from Lin^−^Sca-1^+^c-Kit^+^ (LSK) and Lin^−^c-Kit^+^ (LK) BM cells. (**A-C**) Force-directed graph embedding of publicly available scRNA-seq data from Dahlin et al 2018 showing spatial distribution of the dendritic cell markers (**A**) CD74, (**B**) H2-AA, and (**C**) H2-EB1.

| Antigen | Fluorophore | Vendor | Number | Titer |
| --- | --- | --- | --- | --- |
| Ly6G | SB600 | ThermoFisher | 63-9668-82 | 1/200 |
| CD45 | SB645 | ThermoFisher | 64-0451-82 | 1/100 |
| CD11b | SB780 | ThermoFisher | 78-0112-82 | 1/100 |
| CCR2 | FITC | R&D Systems | FAB5538F-100 | 1/100 |
| CD133 | PercP-eFluor710 | ThermoFisher | 46-1331-82 | 1/100 |
| Ly6C | APC | ThermoFisher | 17-5932-52 | 1/200 |
| Flk-1 | AF700 | ThermoFisher | 56-5821-81 | 1/100 |
| Viability dye | eFluor506 | ThermoFisher | 65-0866-14 | 1/200 |

**Supplemental Table 2**. Panel of antibodies used to determine myeloid cells on bone marrow and blood. For preparation of antibody mixes, Brilliant Stain buffer was used and samples were incubated for 30 minutes at 4^o^C in the dark, followed by two washes. Titers refer to final dilution factors, and staining volume per sample was 100 µl. SB: Super Bright, BV: Brilliant Violet, BB: Brilliant Blue, AF: AlexaFluor, PE: Phycoerythrin, APC: Allophycocyanin.

| Antigen | Fluorophore | Vendor | Number | Titer |
| --- | --- | --- | --- | --- |
| c-Kit (CD117) | SB436 | ThermoFisher | 48-1171-82 | 1/100 |
| FcyRII/III | SB600 | ThermoFisher | 63-0161-82 | 1/100 |
| Sca-1 | SB645 | ThermoFisher | 64-5981-82 | 1/100 |
| Lineage cocktail | FITC | ThermoFisher | 22-7778-72 | 1/50 |
| CD34 | PE | ThermoFisher | MA5-17831 | 1/100 |
| Flk2/Flt3 | APC | ThermoFisher | 17-1357-41 | 1/100 |
| CD127 (IL7Ra) | APC-Cy7 | ThermoFisher | 47-1271-82 | 1/100 |
| Viability dye | eFluor506 | ThermoFisher | 65-0866-14 | 1/200 |

**Supplemental Table 3**. Panel of antibodies used to determine precursor cells on bone marrow. For preparation of antibody cocktail, Brilliant Stain buffer was used and samples were incubated for 30 minutes at 4^o^C in the dark, followed by two washes. Titers refer to final dilution factors, and staining volume per sample was 100 µl. SB: Super Bright, BV: Brilliant Violet, BB: Brilliant Blue, AF: AlexaFluor, PE: Phycoerythrin, APC: Allophycocyanin.

| Antigen | Fluorophore | Vendor | Number | Titer |
| --- | --- | --- | --- | --- |
| F4/80 | SB436 | ThermoFisher | 48-4801-82 | 1/100 |
| Ly6G | SB600 | ThermoFisher | 63-9668-82 | 1/200 |
| CD45 | SB645 | ThermoFisher | 64-0451-82 | 1/100 |
| CD11b | SB780 | ThermoFisher | 78-0112-82 | 1/100 |
| CCR2 | FITC | R&D Systems | FAB5538F-100 | 1/100 |
| CD133 | PercP-eFluor710 | ThermoFisher | 46-1331-82 | 1/100 |
| CD206 | APC | ThermoFisher | 17-5932-52 | 1/100 |
| Flk-1 | AF700 | ThermoFisher | 56-5821-81 | 1/100 |
| Viability dye | eFluor506 | ThermoFisher | 65-0866-14 | 1/200 |

**Supplemental Table 4**. Panel of antibodies used to stain isolated cells from retina. For preparation of antibody cocktail, Brilliant Stain buffer was used and samples were incubated 30 minutes at 4^o^C in the dark, followed by two washes. Titers refer to final dilution factors, and staining volume per sample was 100 µl. SB: Super Bright, BV: Brilliant Violet, BB: Brilliant Blue, AF: AlexaFluor, PE: Phycoerythrin, APC: Allophycocyanin.
